# Lineage-specific retrotransposon co-option reveals a conserved ASH2L isoform as an epigenetic primer of developmental gene promoters

**DOI:** 10.64898/2026.02.25.707914

**Authors:** Eleanor Elgood Hunt, Claudia Vivori, Richard Mitter, Jenie Hannah Johnkingsly Jebaraj, Vedis Agnadottir, M. Joaquina Delás, Eduardo Serna Morales, Thomas Frith, Mark Skehel, Alberto Elosegui Artola, James Briscoe, Folkert J. van Werven

## Abstract

How pluripotent stem cells pre-configure the chromatin landscape to enable timely activation of developmental gene programs remains a central question in developmental biology. Here we demonstrate that the core chromatin regulator ASH2L undergoes a developmentally regulated transcription start site (TSS) switch that generates two protein isoforms with distinct developmental functions: a full-length ASH2L containing an N-terminal intrinsically disordered region (IDR), and a conserved, truncated protein isoform lacking this region. In mouse pluripotent stem cells, a lineage-specific ERV-K retrotransposon drives the upstream TSS, which suppresses the downstream TSS through transcription-coupled SETD2-mediated histone H3 lysine 36 methylation. The resulting stem cell–specific truncated ASH2L assembles into COMPASS complexes and enhances histone H3 lysine 4 trimethylation at promoters governing early development. Despite being restricted to pluripotent stem cells, truncated ASH2L is required for proper axial patterning in gastruloids and for the transition of neural progenitors to motor neurons. This temporal decoupling, wherein a chromatin state established during pluripotency executes its function only after the establishing isoform disappears, defines an epigenetic priming mechanism for developmental timing. Thus, lineage-specific retrotransposon co-option reveals how a dedicated protein isoform configures the pluripotent epigenome for developmental transitions after its own expression has ceased.

## Introduction

The precise temporal and spatial regulation of gene expression is central to the establishment and maintenance of cellular identity during development. One key layer of this regulation is the selection of transcription start sites (TSSs), which can diversify the transcriptome and proteome by generating mRNA isoforms with distinct regulatory or coding potential. Genome-wide studies have revealed that a substantial proportion of mammalian genes initiate transcription from multiple alternative TSSs (Carninci et al. 2006; Makhnovskii et al. 2022; Wang et al. 2008, 2016). Despite their prevalence and developmental regulation, the mechanisms that govern alternative TSS usage, and the functional relevance of the resulting isoforms, remain poorly understood.

Alternative TSSs can contribute to differential gene expression regulation in several ways. They can produce mRNA isoforms that differ in their 5′ untranslated regions (UTRs), thereby influencing mRNA stability, localization, or translational efficiency(Rojas-Duran and Gilbert 2012; Johnstone et al. 2016; Ushijima et al. 2017). In other cases, alternative TSS usage results in transcripts with different coding sequences, leading to the production of protein isoforms with altered domains or cellular functions(Higdon et al. 2024; Chia et al. 2021). These protein isoforms may vary in their subcellular localization, ability to interact with cofactors, or capacity to participate in signaling pathways. The use of alternative TSSs is tightly regulated across tissues and developmental stages, suggesting a functional requirement for dynamic TSS switching.

Studies in yeast and mammalian cells show that alternative TSS usage plays an important role in gene regulation. In yeast, alternative TSSs often generate long undecodable transcript isoforms (LUTIs) containing upstream open reading frames that block translation, and transcription of LUTIs can interfere with transcription from the canonical TSS, leading to the repression of gene expression(Chen et al. 2017; Chia et al. 2017; Cheng et al. 2018; Tresenrider et al. 2021). LUTI-mediated regulation is critical for meiosis and stress responses(Su et al. 2024; Van Dalfsen et al. 2018). In mammals, alternative TSS usage is widespread, but its functional consequences are poorly characterized, with only a few well-defined examples, such as the developmental regulation of Cdk2 isoforms in mice and the MDM2 locus in human cells(Hollerer et al. 2019; Modzelewski et al. 2021).

Mechanistically, the regulation of alternative TSS usage is mediated by transcription and chromatin factors. Transcription from an upstream TSS can repress downstream TSSs through transcription-coupled chromatin changes. For LUTI-mediated and alternative TSS regulation, chromatin factors such as Set2-mediated histone H3 lysine 36 methylation, FACT, Swi/Snf and topoisomerases contribute to the correct TSS balance in yeast(Chia et al. 2021; Tresenrider et al. 2021; Morse et al. 2024). In mammalian cells, chromatin factors such as SETD2 have been implicated in regulating TSS usage(Neri et al. 2017; González-Rodríguez et al. 2022).

Here, using a time course of mESC differentiation into spinal cord motor neurons, we identify a striking example of developmentally regulated TSS switching at the *Ash2l* locus. ASH2L is a core and rate-limiting component of COMPASS and COMPASS-like complexes, which catalyse histone H3 lysine 4 methylation and play essential roles in development and oncogenesis(Shilatifard 2012). We show that in mouse pluripotent stem cells, a lineage-specific ERV-K retrotransposon drives transcription from an upstream TSS, suppressing the downstream canonical TSS through transcription-coupled SETD2-mediated H3K36 methylation. This switch generates a conserved truncated ASH2L isoform that lacks an N-terminal intrinsically disordered region (IDR). This variant enhances H3K4me3 at promoters involved in early development. Although the truncated ASH2L expression is confined to pluripotency, phenotypic defects in axial patterning and motor neuron maturation manifest after the isoform is extinguished. Our findings illustrate how lineage-specific retrotransposon co-option can serve as an evolutionary probe to expose an epigenetic program otherwise obscured by more diffuse regulation across mammalian lineages.

## Results

### Time course of mESCs to motor neuron differentiation identifies alternative TSS switching at the *Ash2l* locus

We performed a time course to determine TSS usage from steady-state RNAs and nascently transcribed RNAs during *in vitro* differentiation of mESCs to spinal cord motor neurons(Gouti et al. 2014; Delás et al. 2023; Rayon et al. 2020). We adopted an established protocol optimised for efficient differentiation in a 7-day time course.

In short, mESCs (cell-line: HM-1) cells were differentiated into spinal cord motor neurons (MNs) through a 7-day time course that transitioned cells from an epiblast-like stem cell (EpiSC) state into neuromesodermal progenitors (NMPs) via bFGF and Wnt activation, followed by patterning into motor neuron progenitors (pMNs) using retinoic acid (RA) and smoothened agonist (SAG), and final maturation into post-mitotic MNs through late-stage Notch inhibition (dibenzazepine, DBZ) (Figure 1A)(Rayon et al. 2020). We validated the differentiation time course through flow cytometry by monitoring cell-type-specific markers (OLIG2, SOX2, ISL1/2, and TUBB3) (Figure 1A and Figure S1A). mESCs transitioned through the EpiSC, NMP, and pMN states to differentiate into ISL1/2/TUBB3 positive motor neurons. We observed 50% of progenitors at day 5 were pMN. The addition of DBZ from day 5 onwards promoted a broader transition into post-mitotic TUBB3-positive mature neurons (Figure S1A).

**Figure 1.**
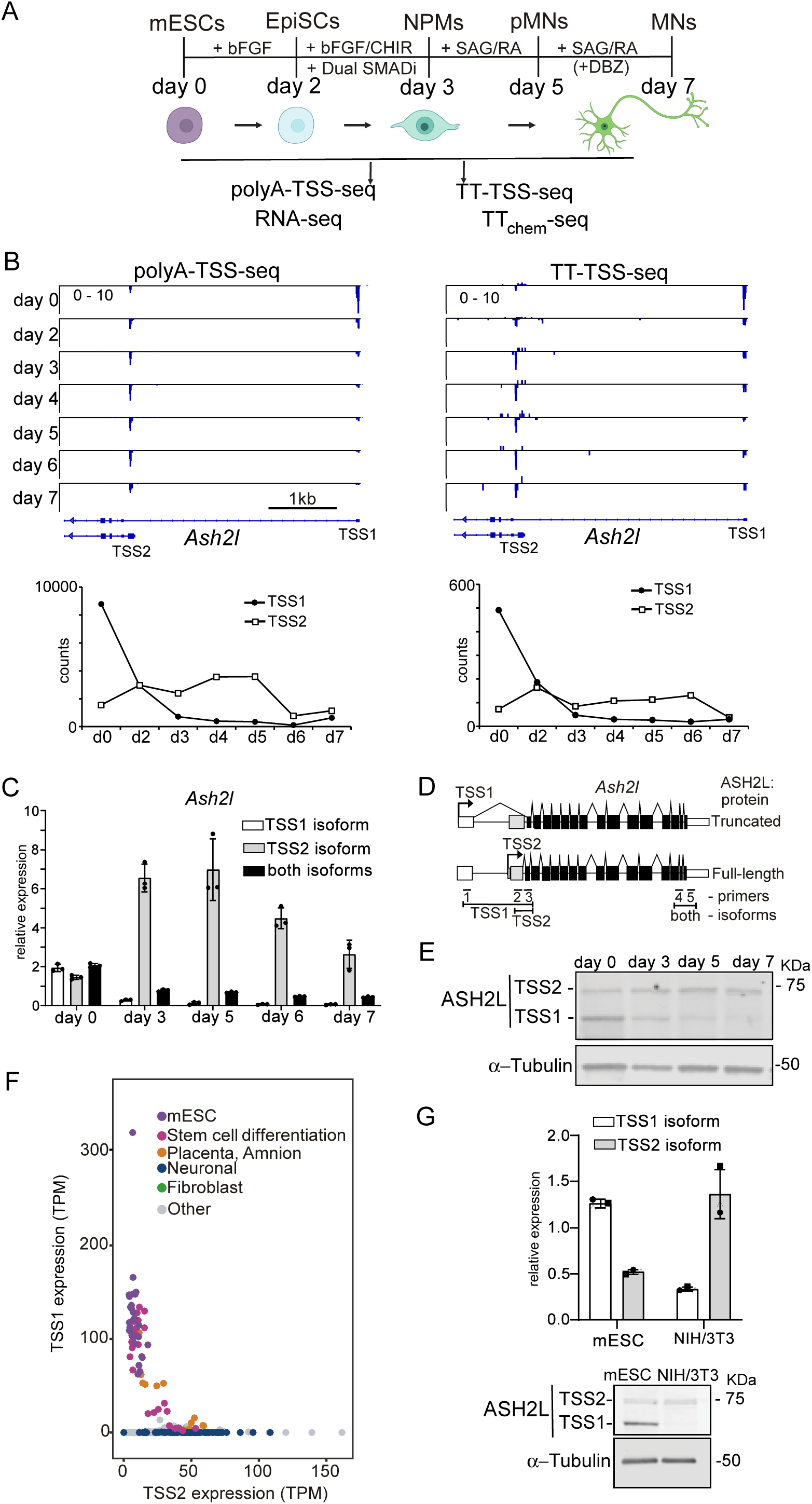
The *Ash2l* locus displays two TSSs that switch in a cell-type-specific manner and encode for different protein isoforms. **(A)** Schematic of the differentiation time course from mouse embryonic stem cells (mESCs) into motor neurons. mESCs first transition through an epiblast-like stem cell (EpiSC, day 2), followed by neuromesodermal progenitors (NMPs, day 3) and motor neuron progenitor (pMN, day 5) stages, and differentiation then into motor neurons (day 7). Overview of transcriptomic assays applied during the time course. Poly(A)-TSS-seq, RNA-seq, TT-TSS-seq, and TT-seq were performed at the indicated time points. **(B)** IGV browser view of the *Ash2l* locus showing data tracks for Poly(A)-TSS-seq and TT-TSS-seq across the time course. Below are shown the quantification of TSS1 and TSS2 usage. The mean signal of N=2 biological repeats is shown. **(C and D)** Quantification of *Ash2l* isoforms by qPCR using primers specific to each isoform. N = 3 biological replicates; mean ± SEM is shown in C. Schematic of the *Ash2l* genomic locus and corresponding protein isoforms detected in D. **(E)** Western blot analysis showing ASH2L isoform expression from TSS1 (truncated protein isoform) and TSS2 (full-length protein isoform) in day 0 (mESCs), day 3, day 5 and day 7 of the time course. **(F)** Tissue-specific expression of TSS1 and TSS2 *Ash2l* usage as determined from published CAGE data. **(G)** Comparison of *Ash2l* TSS1 and TSS2 expression in mESCs and NIH/3T3 fibroblasts. RNA levels (top) are shown as mean ± SD from n=3 biological replicates. Western blot (bottom) probed with antibodies against ASH2L. α−Tubulin is shown as a loading control. A representative experiment of n=3 biological replicates is shown.

Over the time course, we monitored TSS usage from steady-state messenger RNAs (mRNAs) (poly(A)-TSS-seq) and nascently transcribed RNAs using transient labelling with 4-thiouridine (4sU) (TT-TSS-seq) combined with an optimized TSS-seq protocol as described previously(Elgood Hunt et al. 2025). In parallel, we also measured mRNA expression and transcription by RNA-seq and TT_chem_-seq. We found pervasive regulation of alternative TSSs, with about 10% of gene promoters differentially expressing alternative TSSs (Figure S1B). Our genome-wide analysis showed that genes harbouring alternative TSSs are significantly enriched for developmental functions, including morphogenesis and multicellularity (Figure S1C). Among the genes with alternative TSSs, the *Ash2l* locus displayed significant differential TSS usage (Figures 1B, S2A, and S2B). ASH2L is a core component of COMPASS and COMPASS-like chromatin-modifying complexes (Ernst and Vakoc 2012; Shilatifard 2012). We reasoned that ASH2L provides an ideal framework for examining how developmentally regulated alternative TSS usage coordinates gene expression. A comprehensive analysis of the broader TSS landscape within this dataset will be presented elsewhere.

We found that *Ash2l* is transcribed from an upstream TSS1 isoform in mESCs, while at other time points, TSS1 usage decreases (Figures 1B, S2A, and S2B). The downstream TSS2 of *Ash2l* showed the opposite usage pattern. It was the least used in mESCs and the most used from day 3 onwards in the time course. We confirmed the expression of TSS1 and TSS2 *Ash2l* mRNA isoforms using primers targeting exons specific to each *Ash2l* isoform, and a primer pair that detects both isoforms (Figure 1C and 1D). We noted that total *Ash2l* expression was decreased during the time course. Consistent with TSS-seq, TSS1 was the most strongly used in mESCs, while TSS2 was induced upon differentiation into the neuronal lineage.

The two TSSs encode for different *Ash2l* mRNA and ASH2L protein (Figure 1D). In mESCs, the TSS1-derived isoform, which lacks the first exon of the TSS2 isoform, generates a truncated ASH2L protein isoform (Figure 1E). The expression of TSS1-induced ASH2L truncated form was reduced on day 3 and at later time points of the time course. Conversely, the TSS2-regulated isoform encodes the full-length ASH2L protein, which was detected in mESCs at lower levels compared to the truncated isoform. The full-length ASH2L isoform expression increased on days 3, 5, and 7. We conclude that two alternative TSSs drive the expression of two distinct ASH2L protein isoforms, which are differentially used in mESCs and cells differentiating into motor neurons.

Next, we investigated whether the expression of *Ash2l* TSS1 and TSS2 isoforms is regulated across other cell types. We analysed published CAGE datasets (Figure 1F)(Noguchi et al. 2017). We found that neurons and fibroblasts expressed the TSS2 *Ash2l* isoform, while stem cells expressed the TSS1 isoforms. We further examined the expression of the *Ash2l* isoforms in NIH/3T3 fibroblast cells (Figure 1G). We found that the *Ash2l* TSS2 mRNA (full-length ASH2L) was strongly expressed in NIH/3T3 fibroblasts, whereas the *Ash2l-derived* TSS1 mRNA (truncated ASH2L) isoform was detectable at low levels. We conclude that the *Ash2l* TSS1 isoform encodes a truncated ASH2L protein isoform that is the most highly expressed in stem cells, where the TSS2 isoform (full-length ASH2L) is the most highly expressed in differentiated cells.

### *Ash2l* transcription from the TSS1 isoform suppresses TSS2 usage

Given that *Ash2l* TSS1 and TSS2 isoforms have opposite transcription patterns, we wondered whether there is a competitive regulatory environment between the two alternative TSSs. Specifically, we reasoned that TSS1 usage suppresses transcription from TSS2. A similar type of regulation has been described involving noncoding transcription or transcription of overlapping isoforms(van Werven et al. 2012; Martens et al. 2004; Chen et al. 2017; Chia et al. 2017; Hollerer et al. 2019; Lin et al. 2018).

To test this mechanism, we took advantage of the expanded genomic architecture of the *Ash2l* TSSs in the mouse genome. In contrast to the human ASH2L locus, where the two TSS are relatively closely spaced, the mouse *Ash2l* locus provides the spatial resolution for CRISPR interference (CRISPRi) to specifically repress transcription of TSS1 and TSS2 isoforms (Figure 3A)(Gilbert et al. 2013). We used a catalytically inactive Cas9 (dCas9) fused to the transcriptional repressor Krüppel-associated box (KRAB) domain and single-guide RNAs (sgRNAs) to target the *Ash2l* TSS1 and TSS2 regions(Gilbert et al. 2013; Horlbeck et al. 2016; Chen et al. 2019). As a control, we used an sgRNA targeting EGFP as described previously (Horlbeck et al. 2016). In mESCs, the CRISPRi-mediated targeting of *Ash2l* TSS1 and TSS2 isoforms led to 4-fold and 3-fold reduction in expression, respectively (Figure 2A). Notably, repression of the TSS1 isoform triggered compensatory derepression of the TSS2 isoform, resulting in more than 4-fold increase in its expression.

**Figure 2.**
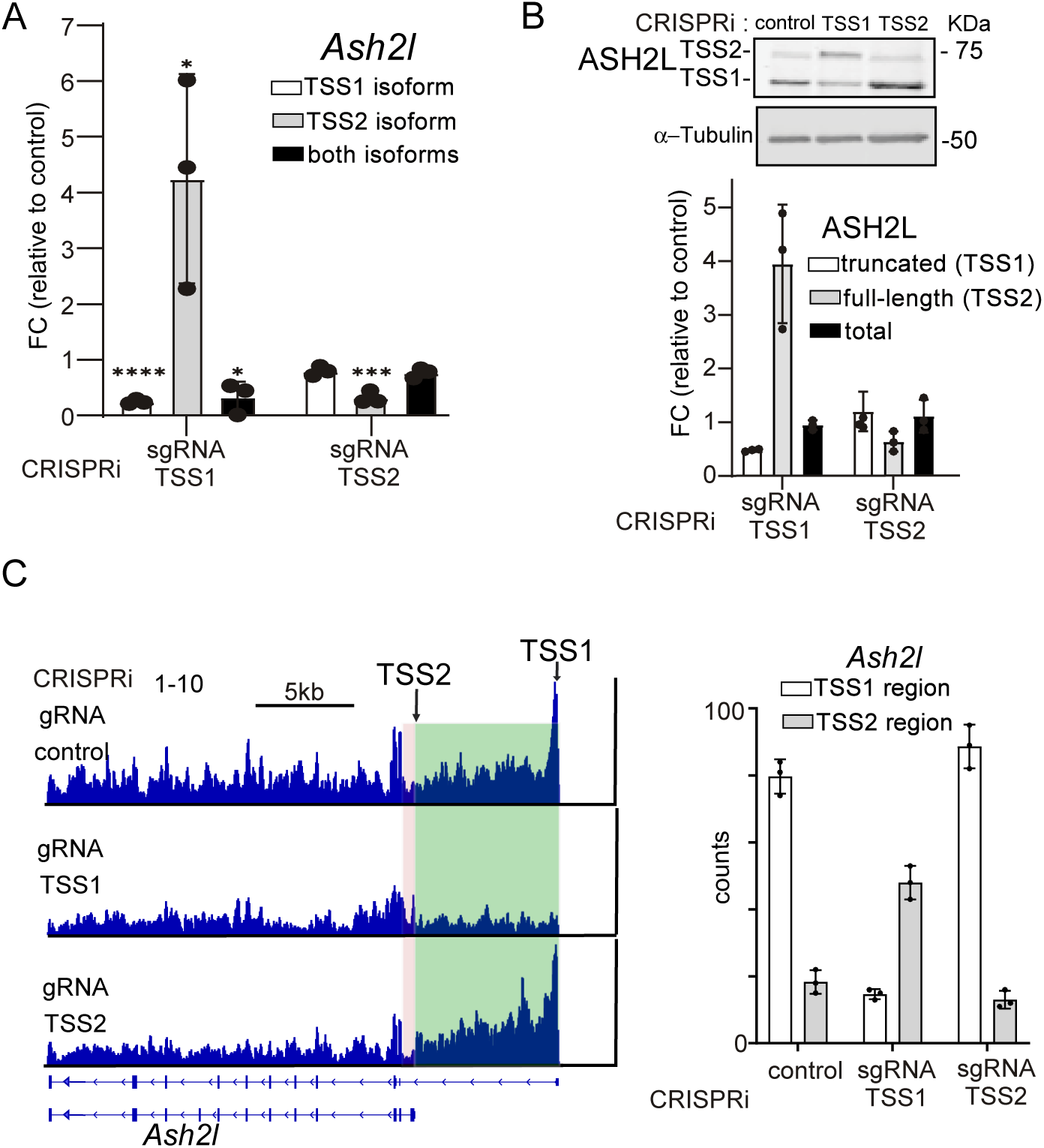
CRISPRi targeting *Ash2l* TSS1 de-represses TSS2 isoform transcription. **(A)** Quantification of total and isoform-specific *Ash2l* expression in cells with CRISPRi targeting either TSS1 or TSS2. mESCs stably expressing dCas9 fused to the KRAB repressive domain were used for the analysis. These cells also expressed sgRNAs targeting either TSS1, TSS2, or a control region. *Ash2l* expression was measured by qPCR using isoform-specific primers and normalised to *Gapdh*. Data are presented as fold change relative to the control. Mean and SD of n=3 biological repeats are shown. Unpaired student’s t-test (* *p*<0.05, ** p<0.01, *** *p*<0.001, **** *p*<0.0001). **(B)** Western blot and quantification of ASH2L protein levels in the CRISPRi mESC cell lines described in A. α−Tubulin was used as a loading control. **(C)** TT_chem_-seq data from CRISPRi mESCs described in A. Highlighted regions indicate TSS1 (green) and TSS2 (red). Quantification of the TSS1 and TSS2 regions is shown on the right. The mean counts are shown for n=3 biological repeats.

**Figure 3.**
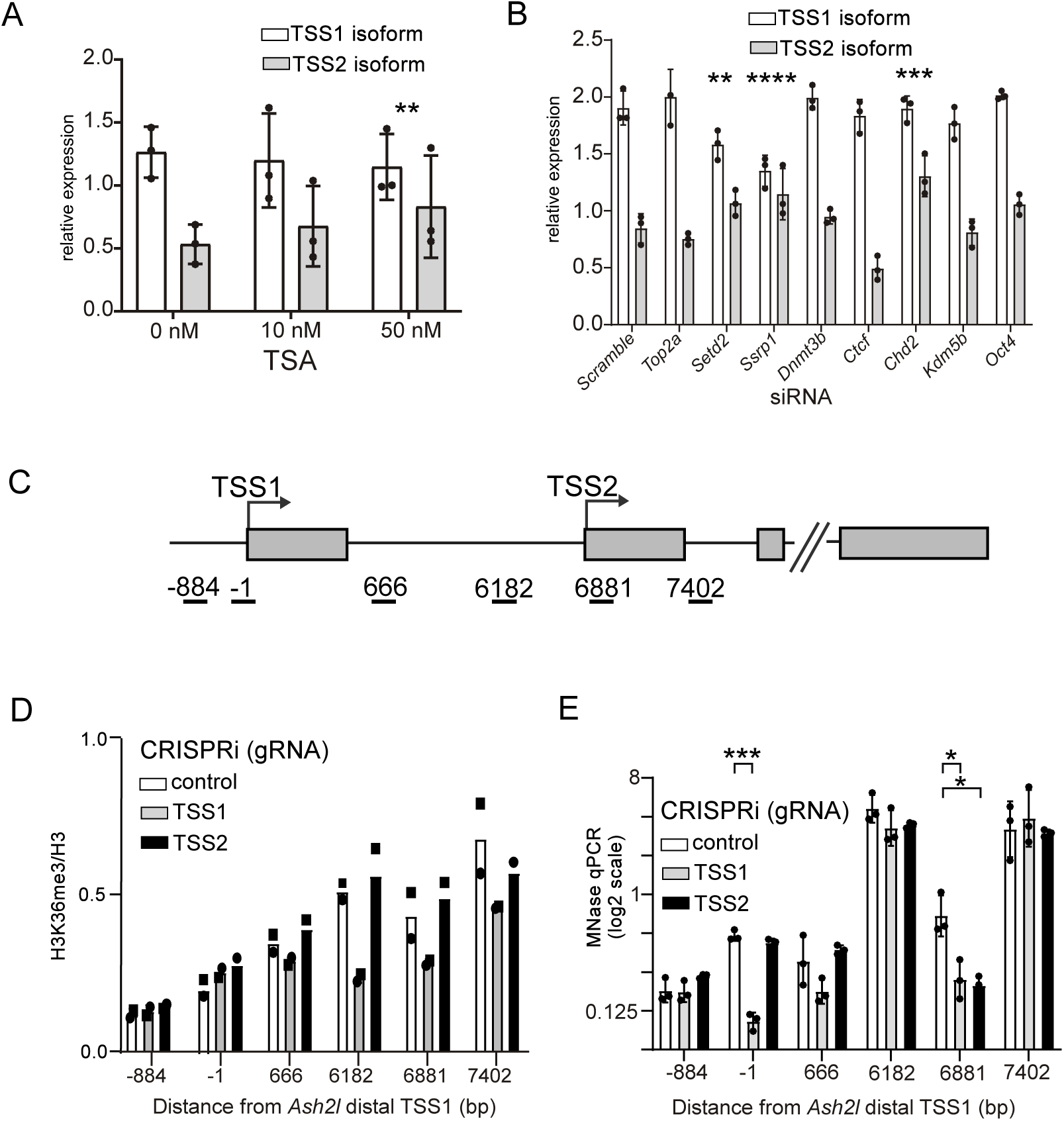
SET2D regulates *Ash2l* TSS1 and TSS2 usage ratio. **(A)** *Ash2l* TSS1 and TSS2 isoform expression following TSA treatment. mESCs were treated for 12 hours before RNA was harvested for RT-qPCR. Expression was normalised to *Gapdh*. Data represent n = 3 biological replicates ± SEM. **(B)** *Ash2l* TSS1 and TSS2 isoform expression in mESCs following knockdown of chromatin or transcription factors using a pool of four small interfering RNAs (siRNAs) per target. Cells were transfected with siRNAs for 48 hours before sample collection. n=3 biological replicates ± SD. Statistically significant pair-wise differences in TSS2/TSS1 isoform ratios compared to the control are indicated (** *p*<0.01, *** *p*<0.001, **** *p*<0.0001). **(C)** Outline of the *Ash2l* locus with primer pairs used for the chromatin immunoprecipitation (ChIP) experiments in D and E. **(D)** ChIP analysis of histone H3 lysine 36 trimethylation (H3K36me3) relative to total histone H3 in control cells and CRISPRi lines targeting TSS1 or TSS2. **(E)** MNase-qPCR analysis of the *Ash2l* TSS1 and TSS2 regions in CRISPRi mESC lines. n=3 biological replicates ± SD are shown.

To test whether the CRISPRi-induced increase in transcription from TSS2 affected protein expression, we examined ASH2L protein isoform levels by western blot analysis. Consistent with observations at the RNA level, CRISPRi targeting TSS1 led to a marked increase in expression of the full-length ASH2L protein isoform (Figure 2B). In NIH/3T3 fibroblasts, where *Ash2l* TSS1 isoform expression is low, we observed a small derepression of TSS2 isoform expression (mRNA, but not protein) when TSS1 was targeted by CRISPRi (Figures S3A and S3B). However, overall ASH2L protein levels did not differ significantly between mESCs and NIH/3T3 fibroblasts. This suggests that the relative ratio of ASH2L isoforms, rather than the total protein pool, is the primary variable regulated by TSS switching across different cell states.

To determine whether the CRISPRi-mediated effects on TSS1 and TSS2 were transcriptional, we performed TT-chem-seq (Figures 2C and S3C). In mESCs, CRISPRi decreased nascent transcription across the targeted TSS1 and TSS2 exons. Again, we observed a reciprocal relationship between the isoforms: transcription of the TSS2 exon was upregulated upon repression of TSS1 isoform transcription, whereas direct targeting of TSS2 resulted in its expected downregulation. In NIH/3T3 fibroblasts, CRISPRi targeting of TSS1 had no detectable effect on transcription from TSS2 (Figure S3C). This may be partly explained by the higher abundance of the TSS2 isoform and the lower abundance of the TSS1 isoform, and the inability of TT_chem_-seq to discriminate between transcript isoforms. Taken together, the analyses suggest that transcription of the TSS1 *Ash2l* isoform suppresses *Ash2l* TSS2 usage.

### SETD2 confers TSS1 mediated suppression of TSS2 *Ash2l* isoform transcription

Previous studies have identified chromatin modifications, such as histone acetylation and methylation, as well as remodelling and topological factors, including FACT and Swi/Snf, as being required for the suppression of gene transcription at overlapping transcription units(Chia et al. 2017; Morse et al. 2024; Tresenrider et al. 2021; Elgood Hunt et al. 2026). We examined whether and which chromatin transcription factors contribute to TSS1 and TSS2 regulation at the *Ash2l* locus.

We first examined whether histone deacetylation is important for maintaining a balanced expression of the two *Ash2l* mRNA isoforms. Histone deacetylation occurs in the wake of transcription to prevent cryptic transcription and to regulate transcription from alternative TSSs(Kim et al. 2012; Carrozza et al. 2005). We treated mESCs with trichostatin A (TSA) to inhibit histone deacetylase (HDAC) activity. As expected, we found that the TSA treatment increased histone H3 lysine 9 acetylation (H3K9ac) (Figure S3D). The increase in H3K9ac correlated with increased *Ash2l* TSS2 usage, while TSS1 usage remained constant (Figure 3A).

Next, we screened eight candidates—including chromatin regulators, topological factors, and transcription factors—to determine how *Ash2l* TSS1 transcription suppresses TSS2 (*Top1a, Setd2, Ssrp1, Dmt3b, Ctcf, Chd2, Kmd5b, Oxt4/Pou5f1*). We knocked down their expression using a pool of four small interfering RNAs (siRNAs) per target in mESCs and NIH/3T3 fibroblasts (Figure S3E and Figure S3F). Except for *Chd2*, the knockdown efficiency of each candidate was at least 50%. We observed that the knockdown of *Setd2* and *Ssrp1* significantly increased the abundance of the TSS2 mRNA isoform (relative to the TSS1 mRNA isoform) in mESCs and NIH/3T3 fibroblasts (Figure 3B and S3F).

SETD2 is the major methyltransferase for histone H3 lysine 36 trimethylation (H3K36me3), and SSRP1 is a component of the FACT complex that regulates nucleosome deposition in the wake of transcription. Both chromatin regulators suppress cryptic transcription and regulate alternative TSS usage in yeast and human cells(Hainer et al. 2011; Carrozza et al. 2005). We decided to focus on SETD2-directed H3K36me3 and determined whether there was evidence of TSS1-dependent H3K36me3 around the TSS2 region. We found that H3K36me3 was present around the TSS2 region in mESCs, but not detectable in NIH/3T3 fibroblasts (which transcribe low levels of the TSS1 isoform) (Figure 3C, 3D and S3G). To determine whether H3K36me3 deposition was dependent on transcription from TSS1, we examined mESCs harbouring CRISPRi. We found that the H3K36me3 signal was significantly reduced in CRISPRi mESCs targeting TSS1, but not when targeting TSS2 or the control (Figure 3D).

SETD2 recruits the well-conserved histone chaperone complex FACT, the de novo DNA methyltransferase DNMT3B, and the H3K4 demethylase KDM5B(Neri et al. 2017; Carvalho et al. 2013; Xie et al. 2011). Collectively, these interactions reinforce a repressive chromatin environment within the gene body, which prevents aberrant transcription initiation. To test this model for the *Ash2l* locus, we assessed nucleosome positioning using micrococcal nuclease digestion combined with quantitative PCR (MNase-qPCR). We found that the nucleosome protection around the TSS2 region was reduced in CRISRPi mESCs targeting TSS1, but not in the control (Figure 3E). Notably, the CRISPRi targeting TSS1 affected nucleosome positioning around TSS1, and CRISPRi targeting TSS2 affected positioning around TSS2, which is consistent with previous reports showing that CRISPR targeting can affect nucleosome positioning (Figure 3E)(Howe et al. 2017b; Isaac et al. 2016).

In summary, our data indicate that *Ash2l* transcription TSS1 induces a repressive chromatin environment by facilitating H3K36me3 and positioning nucleosomes in the TSS2 region. As a result, *Ash2l* TSS1 suppresses the transcription from the TSS2.

### *Ash2l* mRNA and protein isoforms conservation across mammalian lineages

The identification of cell-type-specific switching between *Ash2l* isoforms raises the question of its evolutionary preservation (Modzelewski et al. 2021). We examined how the alternative TSSs and protein isoforms expressed from the *Ash2l* locus are regulated across mammalian species.

First, we assessed whether the N-terminal region of the ASH2L full-length protein sequence (TSS2 mRNA isoform) is conserved. A high degree of similarity in this region was observed across mammalian lineages, with over 75% of residues matching (Figure 4A). The 89-residue region altered between the two mouse ASH2L protein isoforms is predicted to encode an intrinsically disordered region (IDR) (Figure 4B). The IDR is absent in non-mammalian species (e.g., *Gallus gallus, Xenopus tropicalis, Danio rerio, Caenorhabditis elegans,* and *Drosophila melanogaster*), suggesting this region emerged specifically within the mammalian lineage (Figure 4A).

**Figure 4.**
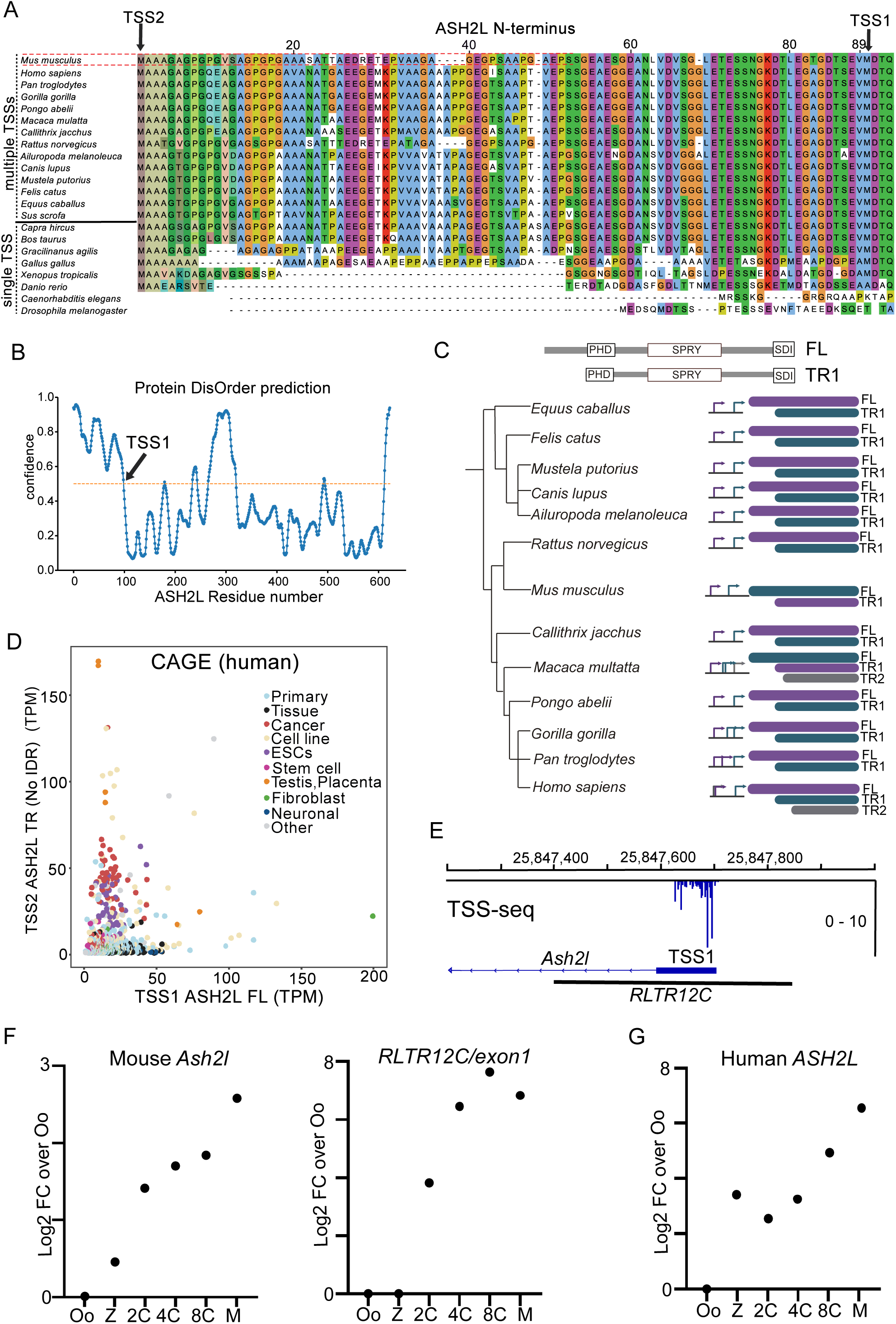
Mouse-specific retrotransposon element rewires *Ash2l* locus TSS usage. **(A)** Protein sequence conservation of the intrinsically disordered region (IDR) at the N-terminus of full-length ASH2L across mammalian species and model organisms. TSS2 (full-length) and TSS1 (truncated) for *Mus musculus* are indicated. Also indicated are species with single or multiple *ASH2L* TSSs. **(B)** IDR confidence scores across the ASH2L protein. Indicated is the TSS1 of *Mus musculus*, where the truncated ASH2L (lacking the IDR) isoform starts. **(C)** Evolutionary tree showing how *Ash2l* transcript and protein isoforms are conserved across mammalian species that express alternative TSSs at the *Ash2l* locus. The arrows indicate the alternative TSSs and are colour-coded to match the ASH2L protein isoforms. Displayed are ASH2L full-length (FL), truncated (No IDR, TR1) and other truncated isoform versions (TR2). **(D)** CAGE expression profiles of human ASH2L TSS1 and TSS2 isoforms. **(E)** IGV snapshot showing TSS-seq signal at the *Ash2l* TSS1 locus and the annotated retrotransposon *RLTR12C* in mESCs. **(F)** RNA-seq data representing the oocyte (Oo), zygote (Z), 2-cell (2C), 4-cell (4C), 8-cell (8C), and morula (M) stages. Shown on the left are fold changes (FC) in *Ash2l* expression, normalized read counts relative to the oocyte (Oo) stage. On the right is displayed the FC for *Ash2l*, specifically at the *RLTR12C Ash2l* retrotransposon junction. **(G)** Similar analysis as in F, except that the ASH2L expression for humans is displayed.

We next investigated the evolutionary conservation of alternative TSS usage in *Ash2l* expression(Modzelewski et al. 2021; Casper et al. 2026). Several mammalian lineages (e.g. *Bos taurus* and *Capra hircus*) only express the full-length ASH2L protein, which includes the conserved IDR (Figure 4A). Conversely, species more closely related to mice (e.g. *Rattus norvegicus, Felis catus* and *Homo sapiens*) use multiple TSSs to generate at least two distinct protein isoforms (Figure 4C). These variants differ by approximately 90 amino acids, effectively mimicking the murine protein isoforms (Figures 4A and 4C). Across these mammalian lineages, the *Ash2l* upstream TSS1 isoform encodes the full-length ASH2L protein, but not the truncated protein, as in mice. *Macaca mulatta* represents a notable exception (Figure 4C), which adopts a tiered alternative TSS architecture: the distal TSS1 encodes a truncated variant lacking about 40 amino acids, whereas the intermediate TSSs drive expression of the full-length isoform, and the proximal TSS generates the conserved truncated protein lacking the first 89 residues.

In humans, ASH2L expression is driven by two TSSs that potentially yield three distinct protein products: a full-length isoform (TSS1), a truncated variant lacking the N-terminal IDR (TSS2), and a shorter isoform that lacks a portion of the PHD domain (TSS1) (Figures 4A, 4C, and S4A). Notably, the upstream TSS1 is predicted to encode the full-length and the most truncated protein isoform. Given that the PHD domain is important for ASH2L function, the most truncated ASH2L isoform is unlikely to be used pervasively. The alternative TSS regulation in humans is less bimodal than we observed in mice, possibly because the alternative TSSs are relatively closely spaced (less than 0.5 kb) (Figure 1F). Nevertheless, TSS1 usage (full-length ASH2) is more dominantly expressed in differentiated tissues, whereas TSS2 (which lacks the N-terminal IDR) is enriched in stem cells and the testis but is depleted in tissues and neuronal lineages (Figures 4D and S4B). Interestingly, the TSS2 isoform is also well expressed in human cancers (Figure S4B). Despite the relatively short distance between the two alternative TSSs in the human genome, our data suggest that the dynamic regulatory logic identified in mice is partially preserved across mammalian lineages, even as the underlying genomic structure has diverged.

In conclusion, whilst the production of two ASH2L protein isoforms, differing in the inclusion of a predicted N-terminal IDR, is conserved, the genomic structure of the TSSs that produce the *Ash2l* mRNA and protein isoforms varies across the mammalian lineages.

### Lineage-specific *RLTR12C* insertion reconfigures the murine *Ash2l* TSS usage

A key contributor to this genomic variation in mammals is the co-option of repetitive elements. Retrotransposable elements frequently hijack host machinery to function as alternative promoters(Modzelewski et al. 2021; Kunarso et al. 2010; Batut et al. 2013). Accordingly, we identified an ERV-K-family long terminal repeat (LTR), specifically of the *RLTR12C* subfamily, that provides the regulatory sequence for the murine *Ash2l* upstream TSS1 mRNA isoform (Figure 4E).

The ERV-K family of retrotransposons are transiently active during early development but silenced at later stages, with high expression in pre-implantation embryos and mESCs, where they can regulate gene expression via stage-specific enhancers and alternative promoters(Göke and Ng 2016; Fu et al. 2019; Oomen et al. 2025). We examined *Ash2l* total and TSS1 isoform expression dynamics during early embryogenesis by analysing published datasets(Boroviak et al. 2018; Kong et al. 2020; Wang et al. 2017; Xue et al. 2013). Consistent with the timing of ERV-K reactivation, *Ash2l* TSS1 exon 1 expression is induced at the 2-cell stage during mouse preimplantation development (Figure 4F).

The mouse-specific *Ash2l* regulatory pattern in early development contrasts with other mammalian lineages (Figures 4G and S4C). In humans, ASH2L expression shows biphasic waves, where ASH2L is induced in the zygote and further induced in the morula stages. In *Macaca mulatta* and *Capra hircus, ASH2L* is relatively unchanged in early embryos, whereas in *Sus scrofa, ASH2L* is induced and peaks at the 8-cell stage (Figures 4G and S4C). Thus, a lineage-specific insertion of ERV-K-family LTR element reconfigured the *Ash2l* locus in mice and is responsible for driving the *Ash2l* TSS1 and TSS2 isoform usage during early development.

### ASH2L isoforms promote histone H3 lysine 4 methylation

ASH2L is an integral subunit of COMPASS and COMPASS-like complexes, which are responsible for the methylation of histone H3 lysine 4(Shilatifard 2012). ASH2L depletion leads to a global reduction in histone H3 lysine 4 methylation, indicating that ASH2L is a rate-limiting component of the COMPASS-like complexes(Ali and Tyagi 2017). How do the two ASH2L protein isoforms control H3 lysine 4 methylation?

We examined the CRISPRi cell lines targeting the TSS1 and TSS2 isoforms. We found that H3 lysine 4 trimethylation (H3K4me3) and, to a lesser extent, di- and monomethylation levels (H3K4me1/2) were diminished in mESCs subjected to CRISPRi targeting either TSS1 and TSS2 (Figure 5A, left panel, Figure S5A). In NIH/3T3 fibroblasts, CRISPRi targeting of TSS2 resulted in a pronounced reduction in H3K4me1/2/3, likely because the TSS2 isoform is the predominantly expressed in these cells (Figure 5A, right panel, Figure S5A). Targeting TSS1 had little effect on H3K4 methylation status in NIH/3T3 fibroblasts.

**Figure 5.**
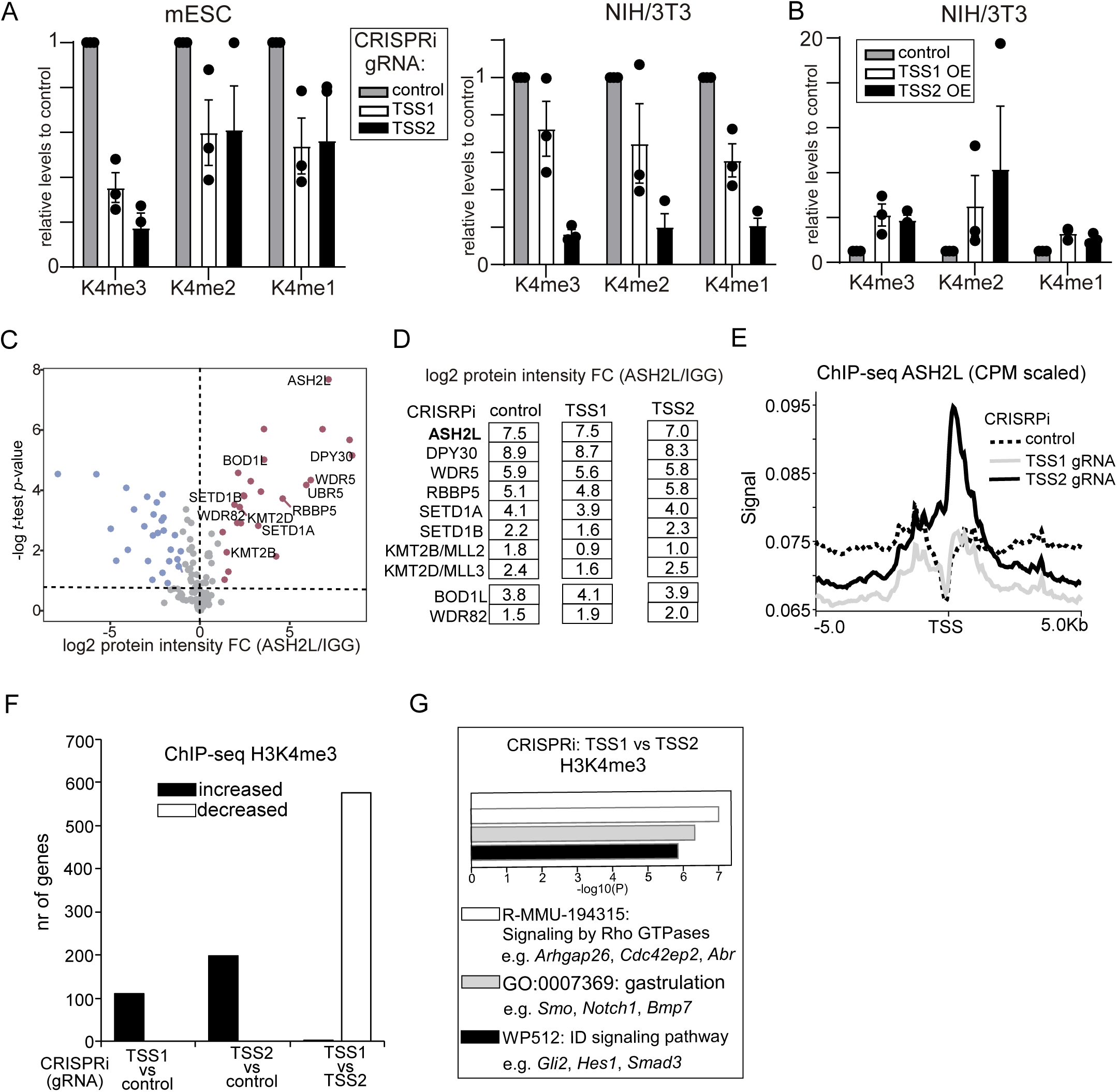
*Ash2l* TSS1 (truncated ASH2L) isoform targets promoters for H3K4me3 important for early development. **(A)** Quantification of histone H3 lysine 4 mono-, di-, and tri-methylation by Western blot in CRISPRi mESCs (left) and NIH/3T3 fibroblasts (right) targeting *Ash2l* TSS1 or TSS2. Data represent the mean ± SD from n = 3 biological repeats. **(B)** Same as (A), except ASH2L (TSS1) truncated and (TSS2) full-length protein isoforms were overexpressed in NIH/3T3 fibroblasts. Data represent the mean ± SD from n = 3 biological repeats. **(C)** Immunoprecipitation followed by mass spectrometry (IP-MS) of ASH2L compared to IgG control in mESCs. **(D)** Same as in C, except comparisons are between control and CRISPRi targeting either TSS1 or TSS2, relative to IgG. Shown are known subunits of COMPASS and COMPASS-like complexes. **(E)** Metagene profiles of ASH2L ChIP-seq at protein-coding genes in control mESCs and CRISPRi targeting TSS1 or TSS2. The signals were centred on TSS. ChIP-seq was normalized by counts per million (CPM). **(F)** ChIP-seq of H3K4me3 comparing control to CRISPRi targeting TSS1 or TSS2 in mESCs. Shown is the number of promoters with significant changes (p < 0.01, fold change > 1.3). **(G)** Gene ontology analysis of promoters with differential changes between CRISPRi targeting TSS1 versus TSS2.

Conversely, we overexpressed either the full-length or the truncated isoform of ASH2L in NIH/3T3 fibroblasts. We observed a marked increase in global H3K4me1/2/3 levels following either full-length or truncated ASH2L overexpression (Figure 5B and Figure S5A). Thus, consistent with ASH2L’s role as a core component of the COMPASS family of histone methyltransferase complexes, both ASH2L protein isoforms act in COMPASS and COMPASS-like complexes to stimulate histone H3 lysine 4 methylation.

### The TSS1–derived ASH2L truncated isoform targets developmentally regulated promoters

Given that TSS1-derived truncated ASH2L isoform expression is limited to mESCs, we reasoned that this variant creates a specialized epigenetic environment at promoters in pluripotent cells. We examined whether the TSS1-derived truncated ASH2L isoform serves this specialized role within COMPASS complexes and regulates histone H3 lysine 4 methylation.

We performed IP-MS in mESCs with control and CRISPRi targeting TSS1 and TSS2. The ASH2L antibody pulled down subunits of both ASH2L isoforms. As expected, CRISPRi targeting TSS1 showed greater enrichment of full-length ASH2L, while targeting TSS2 enriched for the truncated ASH2L isoform (Figure S5B). We observed strong enrichment of the bait protein, ASHL2, along with DPY30, WDR5, and RBBP5—core components that together form the WRAD module, which is essential for the function of COMPASS and COMPASS-like complexes (Figure 5C and 5D)(Ali and Tyagi 2017). In CRISPRi cells, there were no significant changes in WRAD and SETD1A, the primary H3K4 methyltransferase. The only notable difference was a modest reduction in the association of KMT2B/MLL3 in CRISPRi cells targeting TSS1. Our data suggest that the CRISPRi perturbations have no significant impact on the composition of ASH2L COMPASS or COMPASS-like complexes.

Next, we performed ChIP-seq to determine the genomic localization of the ASH2L isoforms and the deposition of H3K4me3 in mESCs with CRISPRi targeting TSS1 versus TSS2. Overall association of ASH2L with chromatin was reduced in mESCs with CRISPRi targeting TSS1 or TSS2 compared to the control when external spike-ins were used for normalisation (Figure S5C). This is consistent with the observation that H3K4me3 levels were reduced in the CRISPRi targeting TSS1 and TSS2 (Figure 5A). When we applied ChIP-seq normalisation by read count, mESCs with CRISRPi targeting TSS2 (thus more dominant expression of the truncated ASH2L protein isoform) showed strong enrichment of ASH2L around the TSSs of coding genes (Figure 5D).

To determine whether the CRISPRi mESCs have altered H3K4me3, we performed H3K4me3 ChIP-seq. As expected, the CRISPRi targeting *Ash2l* TSS1 and TSS2 affected H3K4me3 status at the *Ash2l* locus, with decreased signal around TSS1 (in sgRNA targeting TSS1) and TSS2 (in sgRNA targeting TSS2) (Figure S5D). Across the genome, we found no large differences in H3K4me3 status between control, TSS1 and TSS2 CRISPRi mESCs (Figure S5E). However, we noted a subset of promoters (approximately 600 promoters) had significantly increased H3K4me3 signal (1.3 fold, *p*<0.05) in the mESCs with CRISPRi targeting TSS2 compared to TSS1 (thus, when the ASH2L truncated form is most dominant, compared to CRISPRi targeting TSS1 mESCs where full-length ASH2L is instead most dominantly expressed) (Figure 5F). Gene ontology (GO) analysis indicated promoters associated with genes in Rho GTPase signalling, gastrulation and signalling pathways were significantly enriched (Figure 5G). BMP, Hedgehog, and Notch signalling genes (*Bmp7*, *Smo*, and *Notch1*) showed reduced H3K4me3 at CRISPRi TSS1 compared to TSS2. We also performed H3K4me3 ChIP-seq in NIH/3T3 cells, and found little difference across the CRISPRi cell lines (Figure S5F). This suggests that in mESCs, the truncated ASH2L (driven by the upstream TSS1) promotes H3K4me3 in promoters important for early development.

Given that bivalent chromatin marks are common at early developmental promoters, we investigated whether Polycomb group proteins regulate the specific set of promoters targeted by truncated ASH2L (Bernstein et al. 2006). Polycomb complexes catalyze the trimethylation of histone H3 at lysine 27 (H3K27me3), which, when found in combination with H3K4me3, defines the bivalent chromatin characteristic of developmentally poised genes(Bernstein et al. 2006; Kumar et al. 2021). We analyzed a published dataset and found that the gene set regulated by truncated ASH2L had reduced H3K27me3 signal compared to controls. Meanwhile, H3K4me3 was increased, suggesting that H3K27me3-mediated bivalency is likely not a feature of promoters regulated by the truncated ASH2L isoform (Figure S5G)(Aljazi et al. 2020).

### The TSS1–derived ASH2L truncated isoform is important for early development

Given that the truncated ASH2L isoform targets developmental promoters, we hypothesized that this ASH2L variant primes the chromatin landscape for developmental transitions. Initial support for this hypothesis came from the characterization of a gene trap insertion situated between TSS1 and TSS2 (Stoller et al. 2010). This insertion selectively ablated TSS1 expression while concurrently upregulating TSS2, a reciprocal regulatory shift that mirrors our CRISPRi observations (Figure 2). The loss of TSS1/truncated ASH2L isoform results in embryonic lethality before stage E8.5, coinciding with the onset of gastrulation. To dissect this phenotype, we first assessed the effects on the onset of exit from pluripotency. We found that *Ash2l* isoform expression remained unchanged, and the downregulation kinetics of *Sox2*, *Oct4*, and *Nanog* were indistinguishable across the CRISPRi mESCs during 72 hours of induction of exit from pluripotency (Figure S6A and Figure S6B). Thus, the mESC-specific truncated ASH2L isoform does not play a key role in exiting pluripotency.

To interrogate the developmental context, we employed a gastruloid formation assay to recapitulate trunk morphogenesis *in vitro* (Figure 6A, and see Materials and Methods for details)(Moris et al. 2020). Following mESC aggregation, we recapitulated post-implantation morphogenesis by pulsing aggregates with the Wnt agonist CHIR99021 (48–72h) to trigger symmetry breaking and primitive streak formation (E5.5–E6.5) (Figure 6A). Subsequent axial elongation through 120h drives germ layer specification and the epithelial-to-mesenchymal transition (EMT), through which epiblast cells lose polarity and gain motility to ingress. The Matrigel embedding provides the extracellular matrix cues necessary to facilitate the assembly of “trunk-like structures,” including a rudimentary neural tube and somatic segments.

**Figure 6.**
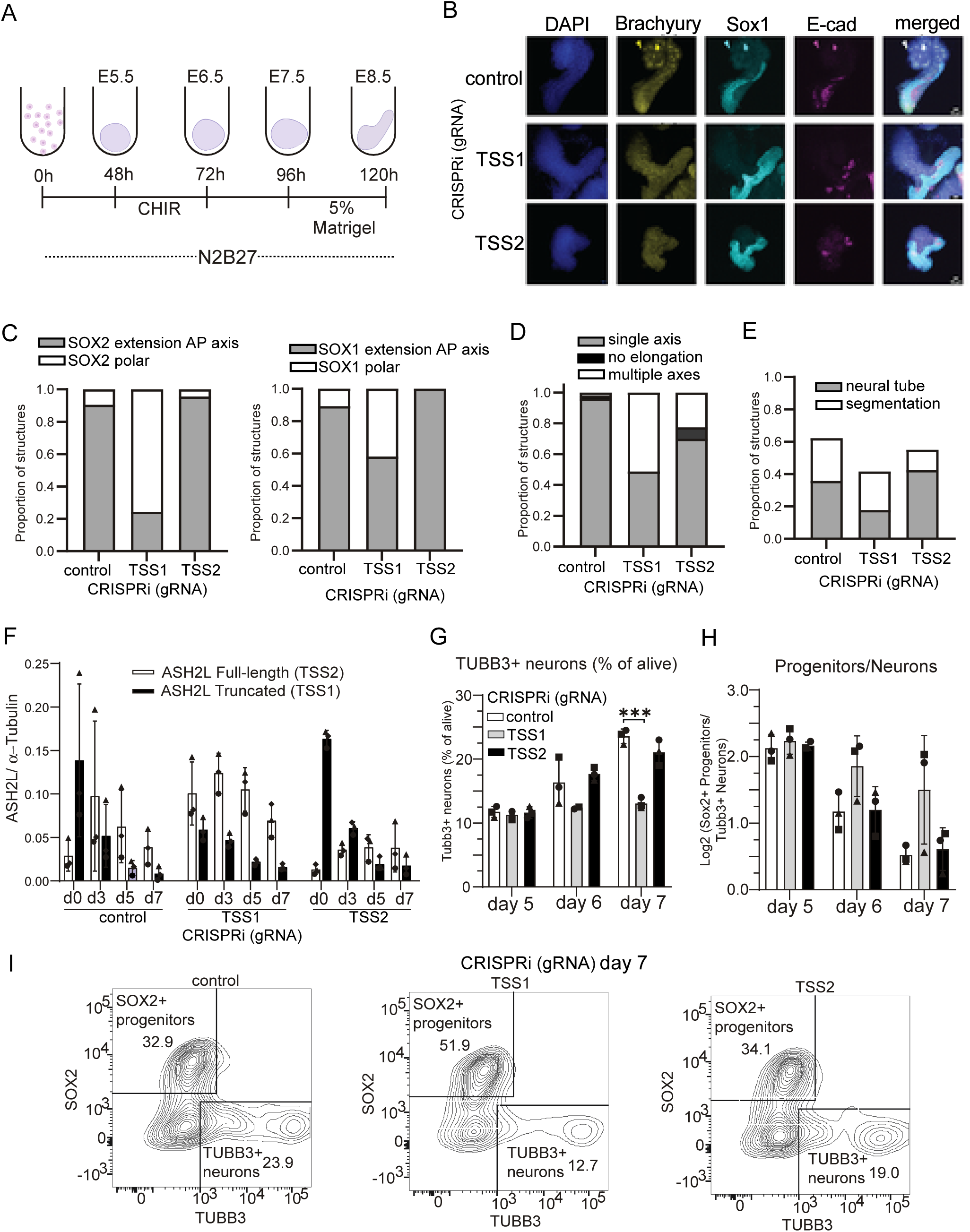
*Ash2l* TSS1 (truncated ASH2L) is important for early development and differentiation into motor neurons. **(A)** Schematic showing the protocol used for generating gastruloids from mESCs as described previously. The time after mESC aggregation is shown, corresponding to the embryonic stage that the gastruloid resembles at each time point. **(B)** Gastruloid assays at 120 hours with Matrigel embedding using mESCs with CRISPRi targeting TSS1 or TSS2. Representative images show staining for SOX1, E-cadherin, Brachyury and DAPI. **(C)** Quantification of SOX2 and SOX1 distribution in control and CRISPRi mESCs targeting TSS1 or TSS2. Gastruloids were classified based on anterior-posterior (AP) distribution and polarity. Shown are the proportions of elongated gastruloids at 120h with 5% Matrigel, with SOX2 staining at the pole or extending along the anterior-posterior (AP) axis (control n=21, TSS1 n=41, TSS2 n=23) and SOX1 staining (control n=28, TSS1 n=43, TSS2 n=15). **(D)** Same experimental setup as C, but quantification of axial morphology. Gastruloids were classified as having a single axis, no elongation, or multiple axes. At least n = 40 gastruloids were quantified per sample. **(E)** Fraction of gastruloids (from A–C) exhibiting segmentation and neural tube-like structures. At least n = 40 gastruloids were quantified per sample. **(F)** Western blot quantification of full-length ASH2L (TSS2 isoform) and truncated ASH2L (TSS1 isoform) during differentiation of mESCs into motor neurons harbouring CRISPRi (TSS1 or TSS2). α−Tubulin was used as a loading control. N = 3 biological replicates; error bars represent SD. **(G)** Flow cytometry quantification of TUBB3-positive neurons in control and CRISPRi targeting TSS1 or TSS2. N = 3 biological replicates; error bars represent SD. **(H)** Same as G, but showing the SOX2+/TUBB3+ cell ratio. **(I)** Representative FACS plots showing TUBB3+ versus SOX2+ cells.

We monitored gastruloid self-organisation in mESCs with CRISPRi targeting TSS1 and TSS2 of *Ash2l*. By 96 hours post-seeding, CRISPRi control and TSS2, Brachyury staining was predominantly localised to the posterior pole, while neural progenitor markers SOX1 and SOX2 were either found proximal to Brachyury or extended along the anterior-posterior axis (Figure S6C and S6D). Notably, SOX17 staining was relatively weak with little evidence of “tube”-like structures forming at this stage. In CRISPRi targeting TSS1, over 40% of gastruloids showed localisation of Brachyury, SOX2, and SOX1 away from the posterior pole in the central region, suggesting that the self-organising capability was disrupted (Figure S6C and S6D).

Also, by 120 hours, control and TSS2-targeted gastruloids exhibited normal axial extension of neural progenitor markers SOX1 and SOX2 (Figures 6B and 6C). Consistent with the 96-hour time point, CRISPRi targeting TSS1 gastruloids frequently failed to break symmetry, maintaining polar SOX1/2 expression (SOX2: 80% vs 10% in control; SOX1: 50% vs 10% in control) and exhibiting a high frequency of ectopic axes (approximately 50%) at 120 hours (Figure 6C and 6D). These multi-polar gastruloids were enlarged, immobile SOX1/2 domains and a significant failure in neural tube formation and segmentation (Figure 6E). While CRISPRi targeting TSS2 caused minor reductions in axial length and E-cadherin expression, it did not disrupt primary patterning (Figures 6B, 6C, 6D, 6E and S6E). Together, these results indicate that the TSS1-derived truncated ASH2L protein isoform is required for proper lineage balance and the coordination of anterior-posterior axial organisation.

### The TSS1–derived ASH2L truncated isoform is essential for motor neuron differentiation

The gastruloid assay provides a readout of anterior-posterior axial elongation and tissue-level organisation in early development(Moris et al. 2020; Veenvliet et al. 2020). The observed axial formation defects in the CRISPRi targeting *Ash2l* TSS1 suggest a potential disruption in the maturation of the neuromesodermal progenitor pool. We hypothesized that failure of axial elongation in gastruloids with CRISPRi targeting *Ash2l* TSS1 impairs motor neuron differentiation.

To investigate this, we directed CRISPRi mESCs targeting TSS1 toward the motor neuron fate (Figures 1A and 6F)(Gouti et al. 2014; Rayon et al. 2020). CRISPRi targeting of TSS1 reduced the truncated ASH2L isoform by ∼3-fold and induced a ∼3-fold increase in full-length ASH2L at day 0, with more subtle differences observed at later time points (days 3, 5, and 7; Figures 6F, S7A, and S7B). Conversely, CRISPRi targeting TSS2 reduced full-length ASH2L levels on days 0 and 3 of the time course.

We quantified stage-specific marker expression to track differentiation kinetics. In control and TSS2-targeted cells, the proportion of post-mitotic TUBB3-positive neurons increased significantly between days 5 and 7. However, CRISPRi cells targeting TSS1 exhibited a ∼50% reduction in TUBB3-positive neurons (Figure 6G). This was accompanied by a sustained SOX2-to-TUBB3 ratio and an accumulation of SOX2-positive progenitors on days 6 and 7, indicating a failure to transition from progenitor to mature neuronal states (Figures 6H, 6I, and S7C).

Importantly, this phenotype was not driven by altered cell viability or defects in early specification, as the expression of motor neuron progenitor (OLIG2) and ventral progenitor (NKX2.2) markers remained largely unaffected (Figure S7D). These data indicate that the TSS1-derived truncated ASH2L protein isoform is required for the maturation of neural progenitors into TUBB3-positive neurons.

### The TSS1–derived ASH2L truncated isoform facilitates developmental gene expression dynamics

How does H3K4me3 established in pluripotent cells shape gene expression dynamics during subsequent differentiation? To investigate, we tracked the expression of truncated ASH2L target promoters (which show enhanced, isoform-specific H3K4me3 in mESCs) throughout a motor neuron differentiation time course (Figure 7A).

**Figure 7.**
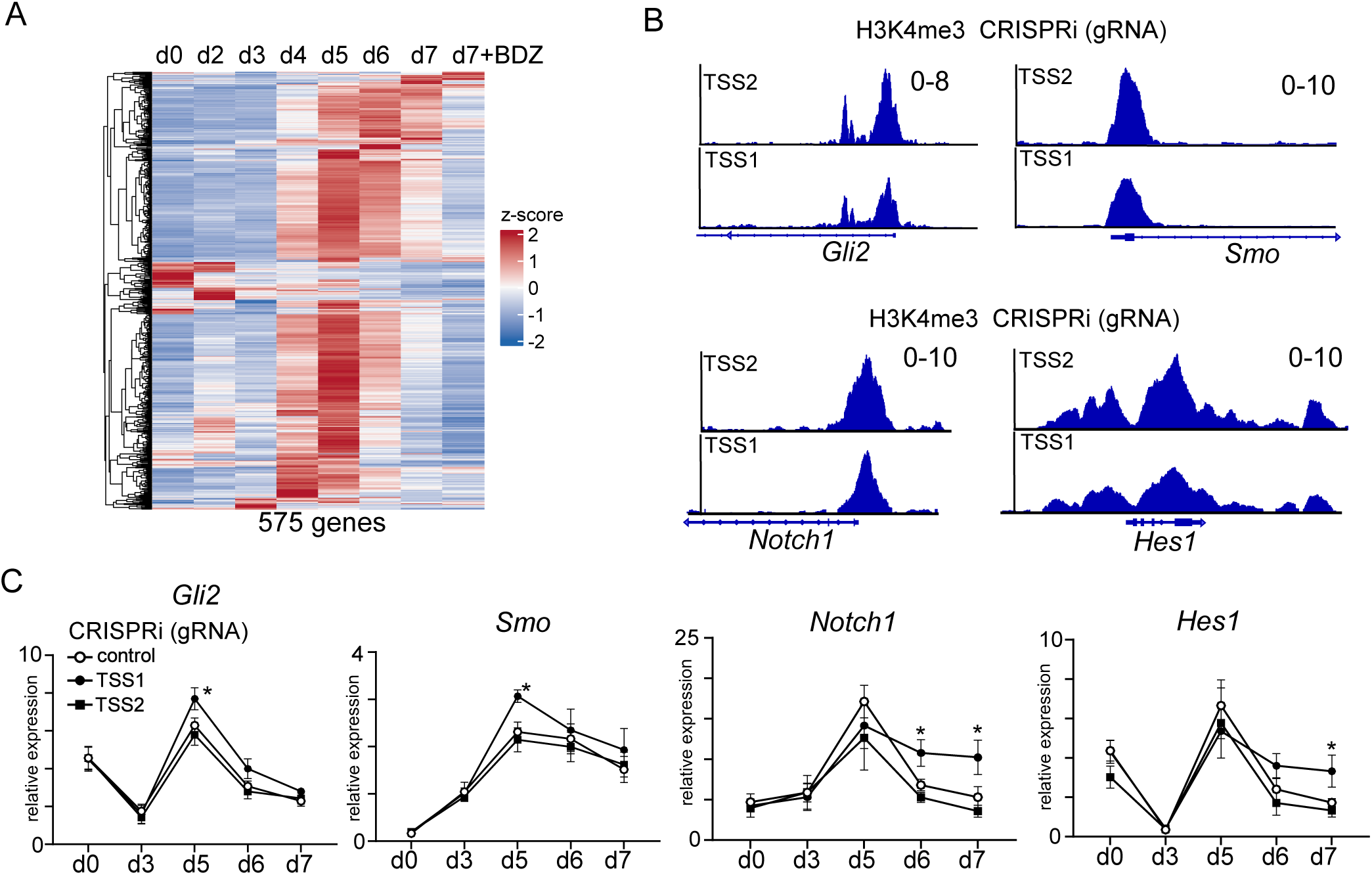
The TSS1–derived ASH2L truncated isoform facilitates developmental gene expression dynamics. **(A)** RNA-seq analysis of time course differentiation of mESCs to motor neurons (see Materials and methods for details). For the analysis, we used 575 genes that show significant differences in H3K4me3 signal as identified in Figure 5F. To compare gene expression levels across the dataset, we calculated z-scores. **(B)** IGV of the *Smo, Gli2, Hes1*, and *Notch1* loci showing H3K4me3 signal in mESCs with CRISPRi (TSS1 or TSS2). **(C)** RT-qPCR measuring the expression of *Smo*, *Gli2, Hes1*, and *Notch1* in the time course of mESCs to motor neurons, comparing control to CRISPRi targeting TSS1 or TSS2. *Gapdh* was used to normalise that data. The mean and SD of n=3 experiments are shown. Statistically significant pair-wise differences (paired *t*-test) compared to the control are indicated (* p<0.05).

The temporal expression analysis across the motor neuron differentiation time course showed that most TSS1-target genes are induced during early differentiation, peaking at the progenitor stage (day 5) before declining as cells transition to mature neurons (days 6–7) (Figure 7A). Among the most dynamically regulated targets were components of the Notch signalling cascade (*Notch1, Hes1, Mib2, Ncstn, Mycn*), the Sonic Hedgehog pathway (*Smo, Gli2*), and transcriptional repressors of the progenitor state (*Tle3, Tle5, Tcf7l1*), pathways that must be precisely timed to promote progenitor expansion and then extinguished to permit neuronal commitment (Imayoshi et al. 2010; Kageyama et al. 2008; Briscoe and Thérond 2013; Torres et al. 2012; Zinin et al. 2014; Muhr et al. 2001; Hirabayashi et al. 2004; Kim and Dorsky 2011).

To validate the functional consequences of altered H3K4me3 in the CRISPRi mESCs, we quantified the expression of four genes (*Smo*, *Gli2, Notch1* and *Hes1*) (Figure 7B). We observed that CRISPRi TSS1-targeted cells exhibited a marked increase in *Smo* and *Gli2* expression compared to controls by day 5, while *Notch1* and *Hes1* expression increased on days 6 and 7 (Figure 7C). This suggests that cells do not fail to activate progenitor-sustaining signals but rather fail to suppress them at the appropriate developmental stage, sustaining a progenitor-identity transcriptional program beyond its proper window.

These findings frame the function of the truncated ASH2L isoform. We propose that it functions as a temporal calibrator by establishing H3K4me3 at progenitor-associated promoters during pluripotency, and it constrains the subsequent dynamics of those genes during differentiation to ensure their timely induction and resolution. The developmental defects caused by depletion of the truncated ASH2L thus reflect a failure of developmental timing.

## Discussion

A central question in developmental biology is how pluripotent cells pre-configure the chromatin landscape to enable the precise temporal activation of developmental gene programs. Here, we demonstrate that a developmentally regulated TSS switch at the *Ash2l* locus provides a mechanistic answer: a stem cell-specific truncated isoform of the core COMPASS subunit ASH2L deposits H3K4me3 at a defined set of developmental gene promoters during pluripotency, establishing a chromatin state that is required for proper axial patterning and timely neural progenitor maturation, yet the isoform itself has been extinguished by the time these developmental events occur. This temporal decoupling between cause and effect defines a form of epigenetic priming that is mechanistically distinct from canonical bivalency(Bernstein et al. 2006). Rather than passively poising genes through co-occupancy of activating and repressive marks, a dedicated isoform of a core chromatin regulator executes a targeted, time-limited preparatory program in pluripotent cells. This model provides a mechanistic basis for the embryonic lethality observed in prior *Ash2l* gene trap studies: the embryo arrests at gastrulation because the epigenetic priming event that should have occurred in the pluripotent epiblast was never completed(Stoller et al. 2010).

We found that in mouse cells, an *RLTR12C* retrotransposon insertion provides the regulatory logic for *Ash2l* alternative TSS usage. Members of the ERV family are frequently active during pluripotency, providing a mechanistic basis for the mESC-specific expression of TSS1(Fu et al. 2019; Oomen et al. 2025). The co-option of the retrotransposon in the mouse has resulted in significantly tighter, cell-type-specific regulation of *Ash2l* TSS usage compared to humans. This regulatory switch uncovered an epigenetic priming function of the ASH2L truncated isoform, which may be more constitutively or subtly managed in other mammalian species. Retrotransposon co-option is a major driver of regulatory innovation, providing a source of alternative promoters that fine-tune essential developmental programs, for example, as demonstrated for the *Cdk2ap1* locus(Modzelewski et al. 2021; Göke and Ng 2016). Our findings at the *Ash2l* locus extend this paradigm by demonstrating that an ERV-K-family LTR exposed a conserved ASH2L isoform function that would otherwise be invisible.

We demonstrate that *Ash2l* TSS1 transcription from retrotransposon element suppresses the downstream TSS2 through transcriptionally coupled SETD2-dependent H3K36 methylation and chromatin compaction. This mirrors paradigms first established in yeast and highlights the conservation of this regulation(Chia et al. 2017; Carrozza et al. 2005). As described in yeast, transcription-directed SETD2-mediated H3K36 methylation modulates chromatin accessibility to prevent downstream initiation. Importantly, this mechanism allows mESCs to drive expression of the ASH2L truncated isoform while maintaining the capacity to derepress the full-length isoform upon differentiation. Thus, the regulatory mechanism described here provides a dynamic means of controlling ASH2L isoform usage.

The stem cell-specific truncated ASH2L isoform, which lacks the N-terminus, retains the ability to assemble into COMPASS and COMPASS-like complexes and to support global H3K4 methylation. Surprisingly, however, we found that this truncated isoform exerts a distinct regulatory role by enhancing H3K4me3 at the promoters of genes involved in gastrulation and early developmental signaling. Throughout a motor neuron differentiation time course, these specific target promoters exhibit tightly regulated temporal expression dynamics, typically peaking at the progenitor stage. The molecular basis for the distinct genomic targeting of the two ASH2L isoforms is an important open question. The N-terminal IDR in the full-length protein is in proximity to the atypical PHD and WH domains of ASH2L that mediate DNA and chromatin contacts, and its removal may increase the accessibility or residence time of these domains at target promoters. IDRs are increasingly recognised as modulators of multivalent chromatin interactions and phase-separated hubs. Thus, the N-terminal IDR of full-length ASH2L may attenuate promoter interactions or reduce the precision of H3K4me3 deposition at the specific developmental promoters targeted by the truncated ASH2L variant. Because ASH2L scaffolds mono-, di-, and trimethylation through distinct COMPASS complexes, the balance of H3K4 methylation states established by the truncated ASH2L isoform likely contributes to regulating the temporal dynamics of target gene expression(Weiner et al. 2012; Howe et al. 2017a). Biophysical characterisation of the N-terminal IDR will be necessary to define the mechanism.

The expanded architecture of the murine *Ash2l* locus, created by the retrotransposon insertion, provided a critical experimental advantage. The retrotransposon restricts the expression of truncated ASH2L to pluripotent stem cells. Additionally, in contrast to human cells, where TSS1 and TSS2 are separated by less than 0.5 kb, the retrotransposon insertion in the mouse *Ash2l* locus provided the necessary spatial resolution for CRISPRi-mediated perturbation, which yielded functional evidence for the distinct regulatory roles of the alternative TSSs. This approach also circumvented the pleiotropic effects of full ASH2L depletion, which disrupts pluripotency exit and would obscure downstream developmental phenotypes(Tsai et al. 2019). CRISPRi targeting *Ash2* TSS1 (truncated ASH2L) significantly impaired gastruloid formation and motor neuron differentiation, which manifested only after the expression of the truncated ASH2L isoform declined, providing direct evidence for the temporal decoupling that defines the priming model.

In summary, this work establishes alternative TSS usage in a core epigenetic regulator as a mechanism for encoding developmental timing information in the chromatin of pluripotent cells. The widespread occurrence of alternative TSSs across the mammalian genome, combined with the frequency of retrotransposon-mediated promoter innovation, suggests that isoform-specific priming functions of the type described here may represent a general strategy by which the genome generates the regulatory precision required for faithful cell fate decisions.

## Methods

### Generation of mESC and NIH/3T3 stable CRISPRi lines

mESCs (HM-1) and NIH/3T3 mouse fibroblast lines were used for the experiments throughout this study. For the expression of catalytically dead Cas9 (dCas9), Plasmid #135448 (Addgene), was used, which contains an ubiquitous Chromatin Opening Element (UCOE) upstream of EF1alpha promoter, and dCas9 fused to a nuclear localisation signal, tagBFP and a KRAB domain (Chen et al. 2019).

For the expression of single-guide RNAs (sgRNAs), Plasmid #84832 (Addgene, CRISPRi V2 plasmid), which contains the mU6 promoter for sgRNA expression, puromycin and BFP selectable markers, was used(Horlbeck et al. 2016). sgRNAs annealed oligo sequences were inserted into Plasmid #84832 into Blp1 and BstXI sites by standard cloning. sgRNAs were designed using CHOPCHOP to be located - 25 to +500 bp, ideally +25 to +100 bp, relative to the TSS (as defined by CAGE FANTOM5 annotations) (Noguchi et al. 2017). A 5’G was appended to each sgRNA to ensure efficient transcription from the murine U6 promoter. Sanger sequencing using a primer located in the U6 promoter was used to confirm cloning of the sgRNA sequence.

Lentiviral infection and sorting were performed as follows. HEK293T/17 cells were plated on gelatin-coated 10-cm dishes and transfected the following day by calcium phosphate precipitation with 10 µg transfer plasmid, 2.5 µg VSV-G (pLP/VSVG), and 7.5 µg pΔ8.9 (pCMV-dR8.91). Viral supernatant was collected 48 and 72 h post-transfection, filtered (0.45 µm), and supplemented with polybrene (8 µg/ml). mESCs in 2i/LIF media, and NIH/3T3 cells in DMEM were seeded, and the media was replaced with virus-containing media.

To obtain stable expression of dCas9 in mESCs and NIH/3T3 fibroblasts, 48 hours after infection, the cells expressing the top 50% of BFP level were sorted by Fluorescence Activated Cell Sorting (FACS). dCas9 expression was confirmed by Western blot, followed by a second sorting, which was carried out after 2-5 passages to ensure purity in dCas9-expressing lines.

Puromycin was used to select cells expressing the sgRNA vector. Optimal concentrations were determined by kill curves in wild-type HM-1 and NIH/3T3 cells, and selection was performed at 0.5 µg/ml (mESCs) or 2 µg/ml (NIH/3T3) starting 48 h post-lentiviral infection. Selection was maintained for at least two passages until non-transduced control cells were eliminated. Oligo sequences and plasmids are in Tables S1 and S2.

### Growth of mESCs and NIH/3T3 cells

mESCs were grown in 2i/LIF (49% DMEM/F12-/- (Gibco), 49% Neurobasal (Gibco), 1X N2 (Gibco), 1X B27 (Gibco), 0.05 mM 2-Mercaptoethanol (Gibco), 2mM L-Glutamine (Gibco), Recombinant mouse 50,000 units LIF (ESGRO), 1 µM PD03259010 (Stemgent), 3 µM CHIR 99021 (TOCRIS), 1% Penicillin/Streptomycin). Cell culture surfaces were coated with 0.15% gelatin in PBS. NIH/3T3 cells were grown in Dulbecco’s Modified Eagle Medium (DMEM, Gibco), 10% Foetal Bovine Serum (FBS, Gibco, Brazil), 1% Penicillin/Streptomycin. All cell lines were maintained at 37°C, 5% CO_2._

### ASH2L isoform cloning and overexpression

ASH2L isoforms were cloned into the pCDNA4/TO mammalian expression vector (Invitrogen) under the CMV promoter. pCDNA4/TO was digested with BamHI-HF and EcoRI-HF (NEB). Total RNA from HM-1 or NIH/3T3 cells was DNase-treated, and poly(A)RNA was purified using Dynabeads Oligo(dT)25 (Thermo Fisher). cDNA was generated using oligo(dT) primers and ProtoScript II Reverse Transcriptase (NEB), followed by RNase H treatment and bead purification. Nested PCR was performed to amplify *Ash2l* isoforms using Q5 or Phusion High-Fidelity DNA Polymerase (NEB), with inner primers adding a C-terminal FLAG or Myc tag. Gibson overhangs were added using KOD DNA polymerase (Merck), and inserts were assembled into pCDNA4/TO using Gibson Assembly (NEB). Constructs were transformed into Stellar competent cells, sequence-verified by Sanger sequencing (Azenta GeneWiz), and transiently transfected as described above. Primer sequences are listed in Table S2.

### Small interfering RNA (siRNA) transfection

siGENOME SMARTpools, containing four siRNA sequences per target (Dharmacon) were used, and the siGENOME Non-Targeting siRNA pool #2 was used for the negative scramble control. siRNAs were dissolved in 1x siRNA buffer (5X: 300 mM KCl, 30 mM HEPES-pH 7.5, 1 mM MgCl2) to a stock concentration of 20 µM. siRNA transfection was carried out as per the manufacturer’s instructions for lipofectamine RNAiMax reverse transfection (Thermo Fisher). Briefly, to transfect a 6-well plate, 4 µl and 7.5 µl lipofectamine per well was used for NIH/3T3 and HM-1 cells respectively, and 10 nM and 25 nM of siRNA respectively. The siRNA and lipofectamine were added separately to 125 µl of Opti-MEM (Thermo Fisher, Reduced-Serum media) each, and then were combined, before the mixture was incubated for 20 minutes. The mixture was then added dropwise to a 1.5 ml cell suspension. Transfection was carried out in antibiotic free media. 24 hours later the media was changed, and samples were collected 48 hours after transfection.

### mESCs to spinal cord motor neurons differentiation

mESCs (HM-1) that had been passaged at least twice post-thawing were used for differentiation. The protocol for spinal cord motor neuron differentiation was adapted from (Delás et al. 2023; Gouti et al. 2014; Rayon et al. 2020). Cells were lifted and resuspended in N2B27 (49% DMEM/F12-/- (Gibco), 49% Neurobasal (Gibco), 1X N2 (Gibco), 1X B27 (Gibco), 0.05 mM 2-Mercaptoethanol (Gibco), 2mM L-Glutamine (Gibco)) with bFGF (10 ng/ml, Bio-Techne, 3139-FB-025) before 85,000 cells were plated on Geltrex- (or Matrigel-) coated 6-well plates in N2B27 with bFGF (10 ng/ml). 48 hours later the media was changed to N2B27 with bFGF (10ng/ml), CHIR99021 (5μM, Axon MedChem, 1386), SB 431542 (10uM, Cambridge Bioscience S0400) and DMH1 (2uM, Cambridge Bioscience, 16679). After 20 hours, the medium was replaced with N2B27 with 100 nM RA (Sigma Aldrich, R2625) and 100 nM SAG (Merck, 566660). After this, the media was replaced every 24 hours for up to 7 days of total differentiation. At day 5, cells were treated with 10 µg/ml notch inhibitor DBZ (Tocris 4889).

For flow cytometry analysis, cells were harvested and counted before being resuspended to 1 million cells per ml in PBS and incubated on ice for 30 minutes with Live/Dead dye (Invitrogen) (1 µl/ml). Cells were then washed once with PBS, resuspended in 50 µl of PBS and fixed for 10 minutes at room temperature with 200 µl 4% PFA. Samples were washed once in PBS and then resuspended in PBS+0.5% BSA. 1 million cells were then taken for staining, and resuspended in 100 µl of PBS with 0.5% BSA and 0.1% Triton-X100 with the primary antibodies either for 2 hours at room temperature or overnight at 4°C (Table S3). UltraComp eBeads (Invitrogen) were used for single stain controls. Cells were then washed and if required, incubated with the secondary antibody in PBS with 0.5% BSA and 0.1% Triton-X100 for 40 minutes at room temperature. Cells were then washed, resuspended in PBS+0.5% BSA, filtered and analysed by flow cytometry.

### Exit from pluripotency

To induce exit from pluripotency, mESCs were cultured in 2i media lacking LIF. Next, cells were plated at a density of 15,000 cells/cm2 in media lacking the two small-molecule kinase inhibitors(Kalkan et al. 2017). Multiple time points were taken over the next 72 hours.

### Flow cytometry

For flow cytometry analysis, cells were harvested and counted before being resuspended in PBS and incubated on ice for 30 minutes with Live/Dead dye (1 µl/ml). Cells were then washed once with PBS, resuspended in PBS and fixed with 4% PFA. Samples were washed once in PBS and then resuspended in PBS+0.5% BSA. Cells were resuspended in PBS 0.5% BSA and 0.1% Triton-X100 with the primary antibodies (Table S3). UltraComp eBeads (Invitrogen) were used as controls. Cells were then washed and, if required, incubated with the secondary antibody in PBS with 0.5% BSA and 0.1% Triton-X100. Cells were then washed, resuspended in PBS+0.5% BSA, filtered and analysed by flow cytometry. Antibodies used are listed in Table S3.

### Gastruloid assay

Gastruloid generation was carried out as described previously(Baillie-Johnson et al. 2015; van den Brink et al. 2020). mESCs (HM-1) were cultured in serum/LIF medium, before being cultured for 24 hours in 2i/LIF medium. Cells were lifted using Accutase, and a single-cell suspension was formed. The cell pellet was washed twice with PBS, before being resuspended in pre-warmed N2B27 (see HM-1 culture protocol). Cells were counted using a haemocytometer for accuracy and diluted to 7.5 cells/µl in pre-warmed N2B27. 40 μl of this suspension was added to the bottom of U-bottomed, non-adherent 96-well plates (Greiner CellStar) using DNA LoBind pipette tips (Elkay). Sides of the plate were tapped to ensure the droplet was at the bottom of the well, and cells were incubated for 48 hours (37°C, 5% CO2). 150 μl N2B27 + 3 μM CHIR99021 was then added to each well, with the pipette tip angled at the bottom of the well to ensure the gastruloid was dislodged from the bottom. After 24 hours, medium was removed and replaced with fresh pre-warmed N2B27 media. At 96 hours, for the gastruloids generated in the absence of Matrigel, this process was repeated. To induce somitogenesis, the media was instead gently replaced with N2B27 + 5% Matrigel (Corning).

Samples were taken at 72, 96 and 120 hours. To each well, media was removed and replaced with 1x PBS, this washing step was performed twice. Fixation was performed with 4% PFA overnight. The gastruloids were then washed three times with 1x PBS. The protocol for gastruloid permeabilisation and antibody staining was previously described(Messal et al. 2021). Gastruloids were permeabilised for four hours at room temperature, on an orbital shaker, with Flash reagent 2 (200 mM borate, pH 7, 250 g/l urea, 80g/l zwittergent). Five washes with PBS 0.2% Triton-X100 (PBT) were next performed, before blocking was performed overnight at 4°C using blocking buffer (PBT, 10% FBS, 0.02% sodium azide, 1% BSA, 5% DMSO). Gastruloids were next incubated with primary antibodies, diluted in the blocking buffer, at 4°C for 72 hours, before three washes in 1x PBS were performed. Incubation with secondary antibodies and DAPI (1:500) was carried out in blocking buffer at 4°C for 24 hours, protected from light, before three washes in 1x PBS were performed. Tissue clearing was carried out using RapiClear 1.49 (SunJin lab), and gastruloids were mounted onto microscope slides in fresh RapiClear.

Images were taken using a Zeiss Invert880 microscope using the Plan-Apochromat 20x/0.8 Ph2 objective or the M27 Plan-Apochromat 10x/0.45 M27 objective (for the 5% Matrigel conditions). Z-slices were taken at 1.6 µm steps. ImageJ was used for image analysis. All images shown in this thesis are maximum intensity Z-projections. Min and max intensity thresholds were set to be equal for all images from that biological repeat. Morphology and the anterior-posterior axis length were measured manually. The mean intensity of each fluorescence channel along the axis was determined. Antibodies used are listed in Table S3.

### RT-qPCR

500 ng of total DNase-treated RNA was reverse transcribed using random primers, Oligo(dT) and Protoscript II (NEB) according to the manufacturer’s protocol. cDNA levels were quantified using PowerUp SYBR Green Master Mix (Life Technologies) and real-time PCR (QuantStudio 7). Normalisation was performed according to *Actin* or *Gapdh* transcript levels. Primer sequences are shown in Table S2.

### Western Blot

For protein extraction, mammalian cells were washed once with PBS and lysed in RIPA buffer with 1x cOmplete Protease Inhibitor Cocktail (Roche), according to the manufacturer’s protocol. The lysate was sonicated for 3 cycles of 30 seconds on/off using a Bioruptor sonicator (Diagenode). Protein concentration was measured using the Pierce Detergent Compatible Bradford Protein Assay (Thermo Fisher).

SDS loading buffer was added (62.5 mM Tris (pH 6.8), 2% β-mercaptoethanol, 10% glycerol, 3% SDS, and 0.017% Bromophenol Blue), and samples were boiled to denature the proteins. 4-20% gradient gels (Bio-Rad TGX) were used for SDS-PAGE (polyacrylamide gel electrophoresis), and proteins were transferred to nitrocellulose membranes using Trans-Blot Turbo Transfer system (Bio-Rad) as per the manufacturer’s instructions. Membranes were blocked with blocking buffer (1% w/v BSA, 1% w/v non-fat powdered milk in PBS with 0.01% v/v Tween-20 (PBST)) for at least 1 hour at room temperature, before incubation overnight at 4°C with primary antibodies in blocking buffer. Membranes were washed with 0.01% PBST, incubated with secondary antibodies in blocking buffer for 1 hour at room temperature, and then washed. Horseradish peroxidase (HRP) signals were detected using enhanced chemiluminescence substrate and exposure to film. IRDye secondary antibodies were detected using an Odyssey Imager (LI-COR).

### ASH2L immunoprecipitation and mass-spectrometry

For co-immunoprecipitation assays, cells were lifted using Accutase or trypsin treatment and washed twice with PBS. The cell pellets were lysed in lysis buffer (50 mM HEPES, 150 mM NaCl, 1mM EDTA, 2.5 mM EGTA, 0.5% Tween, 1x Protease Inhibitor Cocktail, 1 mM PMSF). Following centrifugation, the supernatant was collected, and pre-clearing was performed with 40 µl of a 1:1 mixture of Protein-A and Protein-G Dynabeads (Thermo Fisher, pre-equilibrated with the lysis buffer). 10% of the supernatant was removed for input, and the remainder was incubated overnight at 4°C on a wheel with the immunoprecipitation antibody. 40 µl of a 1:1 mixture of Protein-A and Protein-G Dynabeads (pre-equilibrated with the lysis buffer) was then added. The beads were then washed 5 times with the lysis buffer. For Western blot analysis, the beads were resuspended in SDS loading buffer, and samples were boiled. For mass spectrometry analysis, the beads were washed 7 times with 50 mM ammonium hydrogen carbonate.

The samples were reduced for 1 hour at 37°C using 5 mM DTT, before undergoing alkylation (10 mM IAA, for 30 minutes in the dark). On-bead digestion was carried out with 0.4 μg/μl trypsin overnight at 37°C, before the reaction was terminated with 10% formic acid. C18 clean-up using EV2018 EVOTIP PURE (Evosep) was performed on 1 µg of each peptide eluate, according to the manufacturer’s protocol. Samples were eluted using 50% acetonitrile and vacuum dried to eliminate organic solvent contaminants. 0.1% formic acid was used to resuspend the dried peptides.

Peptide analysis was carried out using nano-scale capillary LC-MS/MS. A flow rate of approximately 300 nl/min was achieved by Ultimate U3000 HPLC (Thermo Fisher). Peptides were captured using a C18 Acclaim PepMap100 5 μm, 75 μm x 20mm nanoViper (Thermo Fisher) before separation was performed on an EASY-Spray PepMap RSLC 2 μm, 100Å, 75 μm x 500 mm nanoViper column (Thermo Fisher). Elution was carried out for 90 mins with an acetonitrile gradient (2-80% v/v). The Orbitrap Eclipse FSN40303 (Thermo Fisher) mass spectrometer was utilised.

Data was blasted against the UniProt mouse proteome. MaxQuant software version 2.0.3.1 was used for analysis, using label-free quantification (LFQ). The minimum peptide length was 7 residues, and the false discovery rate (FDR) threshold was 0.01. Perseus 1.6.14 was used for subsequent analysis. Potential contaminants, reverse hits, and hits lacking unique peptides were removed. LFQ intensities were log-transformed and compared between samples using a permutation-based FDR *t*-test. An FDR threshold of 0.05 and a fold-change threshold of 1.5 were utilised for testing for significantly different interactions. To compare the ASH2L immunoprecipitation and the immunoglobulin G (IgG) immunoprecipitation, missing value imputation was carried out, whereby missing values were replaced with random numbers taken from a normal distribution. The normal distribution was calculated per sample with a 1.8 standard deviation down shift and a width of 0.3.

### RNA isolation and 4sU labelling

RNA was extracted from mESCs and NIH/3T3 fibroblasts by the standard TRIzol extraction procedure (Thermo Fisher). rDNase (Qiagen) digestion was carried out, and purification was performed by spin column (Qiagen) or by phenol: chloroform extraction and ethanol precipitation.

4sU labelling was carried out as detailed previously (Gregersen et al. 2020), with a few modifications. Cell culture media was removed, filtered, and 1 mM 4sU (Glentham Life Sciences) was added. The cells were returned to the incubator and labelled with 4SU for 15 minutes before medium aspiration and TRIzol addition. 1/5 volume of chloroform was added, followed by isopropanol precipitation. The pellet was then washed with 85% ethanol, dried, and resuspended.

RNA integrity was checked for a selection of samples using Bioanalyzer (Agilent Technologies). 4sU incorporation was assessed using a slot blot as described previously(Elgood Hunt et al. 2025). RNA biotinylation and purification of the biotinylated RNA were carried out as described previously (Gregersen et al. 2020; Elgood Hunt et al. 2025).

### TT-TSS-seq and Poly(A)-TSS-seq

The TT-TSS-seq and Poly(A)-TSS-seq methods were previously described in detail(Elgood Hunt et al. 2025). In short, total RNA was dephosphorylated with QuickCIP (NEB, 1.2 U/µg RNA. RNA was extracted using phenol/chloroform/isoamyl alcohol. RNA was treated with mRNA decapping enzyme (NEB, 0.7 U/µg RNA) followed by purification by phenol/chloroform/isoamyl alcohol. 5’ adaptor ligation was performed with T4 RNA ligase I (30 U). RNAClean XP (1.8X ratio, Beckman Coulter) was used to remove unligated adaptors, according to the manufacturer’s instructions. Poly(A)+ RNA was enriched using Dynabeads Oligo(dT)25 (Thermo Fisher), or 4sU+ RNA was purified as detailed in the TT_chem_-seq protocol (Gregersen et al. 2020). RNA was fragmented, and 3’ ends were fixed by QuickCIP treatment (NEB, 30 U). Pre-adenylated 3’ adaptor (2 µM) ligation was carried out by T4 RNA ligase 2, truncated KQ (NEB, 200 U). Reverse transcription was carried out by SuperScript III (2 µl, Invitrogen). AMPure XP beads (Agencourt, Beckman Coulter) were used to clean up the cDNA. cDNA pre-amplification was performed with i5_s and i7_s (300 nM each) primers and Phusion HF PCR Mastermix (Thermo Fisher). 6 PCR cycles were carried out. ProNex beads (Promega) with a 1:2.95 ratio to the sample were used to remove sequences less than 55 nt (including primer dimers). Final amplification was carried out with NEBNext i50 and i70 primers (500 nM each) and Phusion HF PCR Master Mix, for 8 cycles. Amplification was checked by using Novex 6% TBE (Tris/Borate/EDTA) gel and staining with SYBR Green. Libraries were sequenced with Illumina NovaSeq 6000 with at least 50 million reads per sample.

### RNA-seq and TT_chem_-seq

For RNA-seq, libraries were prepared using NEB Ultra II, with poly(A)+ purification, according to the manufacturer’s protocol. Sequencing was performed on NovaSeq 6000 (Illumina) using 100 bp paired-end reads, to at least 50 million reads.

For TT_chem_-seq, at least 10 ng of eluted RNA was used for library preparation with NEBNext Ultra II Directional Poly(A) mRNA, according to the manufacturer’s protocol for fragmented RNA. Libraries were sequenced using NovaSeq 6000 (Illumina) as 100 bp paired-end reads, to at least 50 million reads.

### ChIP and ChIP-seq

The chromatin immunoprecipitation (ChIP) protocol was adapted from (Esnault et al. 2014).1.6 million mESCs in 2i/LIF media and 2.5 million NIH/3T3 cells in DMEM were plated in 15-cm plates. After 48 hours, the cells were fixed for 10 minutes at 37°C with a 1% formaldehyde solution in PBS, before the reaction was quenched with 250 mM glycine. Cells were washed twice in cold PBS before being collected using a cell scraper in 3 ml of cold PBS containing 1x Protease Inhibitor Cocktail (Roche). Following centrifugation the cell pellet was lysed in 5 ml of lysis buffer (5 mM HEPES pH 8.0, 85 mM KCl, 0.5% Triton-X-100, 1x Protease Inhibitor Cocktail, 1 mM PMSF), then washed in 5 ml of buffer (5 mM HEPES pH 8.0, 85 mM KCl, 1x protease inhibitor cocktail, 1 mM PMSF), before being resuspended in buffer (50 mM Tris-HCl pH 8.0, 1% SDS, 10 mM EDTA, 1x Protease Inhibitor Cocktail, 1 mM PMSF) and snap-frozen on dry ice. Once the samples were thawed, sonication was carried out with 5 cycles of 30 seconds on/off (Bioruptor Plus (Diagenode), low intensity setting). Following centrifugation, 10% of the sample was removed for input. The input sample was treated with proteinase K (NEB) overnight at 65°C and purified using Zymo ChIP DNA Clean & Concentrator, and used to check for sonication efficiency. The remaining sample was precleared with a 1:1 mix of protein-A and protein-G Dynabeads (pre-equilibrated in PBS 0.1% BSA) for 1 hour at 4°C. The pre-cleared sample was then mixed with the antibody overnight at 4°C, before 1:1 mix of protein-A and protein-G Dynabeads (pre-equilibrated in PBS 0.1% BSA) were added for 2 hours. The beads were then washed 3 times in FA/SDS solution (50 mM HEPES KOH pH 7.5, 150 mM NaCl, 1% Triton-X-100, 0.1% Na deoxycholate, 1x Protease inhibitor cocktail, 1 mM PMSF) with the last wash being performed for 10 minutes at 4°C. The beads were then washed once in WB buffer (10 mM Tris-HCl pH 8, 0.25 M LiCl, 1 mM EDTA, 0.5% NP40, 0.5% Na-deoxycholate), once in TE (Tris/EDTA) buffer pH 8.0, before being eluted in elution buffer (25 mM Tris-HCl pH 7.5, 5 mM EDTA, 0.5% SDS) at 65°C for 25 minutes. The samples were treated with RNase (NEB) for 30 minutes at 37°C before being reverse crosslinked overnight with Proteinase K at 65°C, and finally purified with Zymo ChIP Clean & Concentrator. For ChIP-seq, libraries were prepared using the NEBNext Ultra II DNA Library Prep Kit for Illumina accordingly. For qPCR, samples were diluted 5-fold, and 1 µl was used per reaction.

The protocol for micrococcal nuclease (MNase) qPCR analysis was adapted from(Yu et al. 2020). Cells were lifted using Accutase or trypsin, the pellet was washed once in PBS and lysed in buffer A (10 mM Tris-HCl, pH 7.5, 60 mM KCl, 15 mM NaCl, 3 mM MgCl_2_; 1 mM DTT, 0.1% NP-40) for 10 minutes on ice. Following centrifugation, cells were resuspended in buffer A without the NP-40, and 10% input was removed. The micrococcal nuclease (NEB) reaction was carried out according to the manufacturer’s protocol (4.5 µl MNase, at 37°C for 12 minutes). Next, DNA was treated with RNase A (NEB) and then treated with 1% SDS and proteinase K (NEB) for 2 hours at 55°C. Phenol-chloroform extraction and ethanol precipitation were carried out. DNA concentration was measured with a Qubit 4 fluorometer (Thermo Fisher) using Qubit dsDNA HS (according to manufacturer’s protocol), and an equal amount of DNA was used for the qPCR reaction. To calculate nucleosome occupancy, we first determined the signal relative to the input and then corrected this for an intergenic region downstream of the *Ash2l* locus.

### Bioinformatic analysis

TT-TSS-seq and Poly(A)-TSS-seq read demultiplexing was performed using Ultraplex with parameters “--phredquality 15 --min_length 0” (Wilkins et al. 2021). Adaptor trimming was performed using Cutadapt with parameters “--minimum-length 20” (Martin 2011). Bowtie 2 was used for pre-mapping to ribosomal and small RNAs (Langmead and Salzberg 2012). Genome mapping was performed using STAR, for mouse samples with Ensembl GRCm38 (mm10) release-89 annotation(Dobin et al. 2013). Umitools was used for deduplication(Smith et al. 2017). BEDTools was used to identify the 5’-most nucleotide (tag) and for genome-wide comparative analyses (Quinlan and Hall 2010).

RNA-seq and TT_chem_-seq time course analysis read processing using nf-core/rnaseq (version 3.13.2) (Ewels et al. 2020). Adaptor and quality trimming were performed using Trimgalore (minimum trimmed read threshold: 10,000). Genome mapping was performed using STAR with Ensembl GRCm38 (mm10) release-89 annotation (minimum uniquely mapped read threshold: 5%)(Dobin et al. 2013). Transcript assembly was performed using StringTie(Pertea et al. 2015).

For the analysis of a public ChIP dataset (GSE157778), H3K27me3 and H3K4me3 for ChIP binding profiles over selected genes. Counts-per-million (CPM) normalised BigWig files representing H3K27me3 and H3K4me3 ChIP binding across the mouse mm9 genome assembly were downloaded from GEO under series GSE157748(Aljazi et al. 2020). DeepTools’ ComputeMatrix function(Ramírez et al. 2016) was used to calculate mean ChIP coverage across the TSS of selected genes +/-1kb, using 10bp bins. Genes were selected from a custom list of putative ASH2L regulators and compared to control sets: i) all protein-coding genes and ii) a random subset of protein-coding genes that did not overlap the custom list. Read-depth profiles and heatmaps were plotted directly from the output matrix using deepTools’ plotHeatmap function(Ramírez et al. 2016). The profiles were collapsed at a metagene level by taking the mean of each bin across all genes. The heatmaps show z-score normalised intensity values per gene.

ChIP-seq of ASH2L and H3K4me3 were processed using the nf-core chipseq pipeline (version 2.0.0) against the composite mouse (GRCm38) and *Drosophila* (BDGP6-46) genomes. Transcript annotations were derived from the mouse Ensembl release 102. For Peak calling, we used MACS2 against matched input controls, and the ENCODE mm10 blacklist was used to remove false positives. BigWig files representing genome-wide coverage were generated directly from the BAM files and scaled based on the *Drosophila* spike-in. A per-sample scale factor was derived by dividing the smallest observed spike-in read count across all samples by the equivalent count from the sample in question. The Bioconductor packages DiffBind and DESeq2 were used to test for differential binding between H3K4me3 conditions. Consensus peaks (e.g. peaks identified consistently in the 3 replicates of each group) were resized to a width of 2kb centred on their summits. Next, the Control (input) read counts were subtracted for each site in each sample before normalisation using the spike-in reads via a “background” approach. Essentially, the spike-in genome was divided into “background” bins of sufficient size (15 kb) so as not to show differential enrichment between samples. Counts over these bins were then used to generate per-sample scale-factors and passed to DESeq2 for significance testing of pairwise comparisons, thresholding for significance at FDR<0.05. A post filter (minimum peak signal to >=300, fold-change>1.25) was used to remove weak peaks after manual inspection.

TT-seq from the CRISPRi experiment was processed using nf-core rnaseq pipeline (version 3.12.0) using RSEM/STAR for abundance estimation against genome version GRCm38 with associated Ensembl release 102 transcript annotations. The “--additional_fasta” flag was set to allow alignment of simultaneously reads against the *Saccharomyces cerevisiae* R64-1-1 genome, permitting the quantification of yeast spike-in reads. Bigwig files representing genome-wide coverage were created directly from the BAM files and normalised to CPM based on total mapped reads for visualisation.

### Statistical analysis

Statistics used for the figures are indicated in the figure legends.

## Data Availability

The accession number for the sequencing data reported in this paper is deposited in GEO under accession numbers GSE320328 and GSE320327. The mass spectrometry proteomics data have been deposited to the ProteomeXchange Consortium via the PRIDE partner repository with the dataset identifier PXD074260.

## Acknowledgements

We thank the Crick Genomics STP for sequencing the libraries and the Flow Cytometry STP for training and access to the FACS instruments. We also thank the members of the van Werven lab for the critical reading of the manuscript.

---

Table S1. Primers used.

Table S2. Plasmids used

Table S3. Antibodies used.

## Figure legends

**Figure S1. TSS-seq and TT-TSS-seq analysis of mESCs differentiating to motor neurons.**

**(A)** FACS analysis of SOX2, OLIG2, ISL1/2, and TUBB3 expression during differentiation. Showing the percentage SOX2+ cells (top left) representing progenitors, OLIG2+ SOX2+ cells (top right) representing motor neuron progenitors, ISL/2 and TUBB3+ cells representing mature motor neurons, and TUBB3+ cells representing mature neurons. **(B)** TSS-seq and TT-TSS-seq analysis. Plotted are the number of TSSs and promoters with alternative TSSs that are differentially used over time. **(C)** Gene ontology analysis of genes with alternative TSS usage, comparing mESCs to day 7 of the motor neuron differentiation time course.

**Figure S2. *Ash2l* TSS1 and TSS2 isoform transcription and expression throughout differentiation from mESCs to motor neurons.**

**(A)** RNA-seq dataset of neuronal differentiation time course from day 0 to day 7. Displayed are the IGV tracks for each time point. TSS1 (green) and TSS2 (red) are highlighted. **(B)** Similar analysis as in A, except that TT_chem_-seq data is displayed.

**Figure S3. SETD2 regulates *Ash2l* TSS1 and TSS2 usage.**

**(A)** Quantification of total and isoform-specific *Ash2l* expression in NIH/3T3 cells with CRISPRi targeting either TSS1 or TSS2. 3T3/NIH fibroblasts expressing dCas9 fused to the KRAB repressive domain were used for the analysis. These cells also expressed sgRNAs targeting either TSS1, TSS2, or a control region. *Ash2l* expression was measured by qPCR using isoform-specific primers and normalised to *Gapdh*. Data are presented as fold change relative to the control. The mean and SD of n=3 biological repeats are shown. Unpaired student’s t-test (* *p*<0.05, ** p<0.01, *** *p*<0.001, **** *p*<0.0001). **(B)** Western blot analysis of ASH2L protein levels in the CRISPRi 3T3/NIH fibroblast lines described in A. α−Tubulin was used as a loading control. **(C)** TT_chem_-seq data from CRISPRi 3T3/NIH fibroblasts described in A. Highlighted regions indicate TSS1 (green) and TSS2-specific regions. Quantification of the TSS1 and TSS2 regions is shown on the right. The mean counts are shown for n=3 biological repeats. **(D)** Western blot using acetylated histone H3 lysine 9 levels after TSA treatment with 0, 10, and 50 nM in growth medium for 12 hours. As a loading control, α−Tubulin is shown. **(E)** siRNA knockdown efficiency of different chromatin and transcription factors in mESCs. mESCs were transfected with a pool of four siRNAs per target, targeting individual chromatin or transcription factors for 48 hours. RNA was extracted and analyzed by RT–PCR. Expression levels were normalized to *Gapdh*, and fold changes were calculated relative to the scrambled siRNA control. Data represent the mean ± SD from n = 3 biological replicates. **(F)** *Ash2l* TSS1 and TSS2 isoform expression (left) in 3T3/NIH fibroblasts following knockdown of chromatin or transcription factors using siRNAs. Cells were transfected with siRNAs for 48 hours before sample collection. Knockdown efficiencies normalised to controls were determined by RT-qPCR (right). n=3 biological replicates ± SD. Statistically significant pair-wise differences in TSS2/TSS1 ratios compared to the control are indicated (** p<0.01). **(G)** ChIP analysis of H3K36me3 relative to total histone H3 in control 3T3/NIH fibroblasts and CRISPRi 3T3/NIH fibroblasts targeting TSS1 or TSS2. n=3 biological replicates ± SD.

**Figure S4. *Ash2l* isoform expression in human and mammalian lineages.**

**(A)** ASH2L locus structure and alternative TSS usage in human cells. Displayed are the CAGE data from human cells. TSS1 can encode for full-length ASH2L and a truncated version lacking the first 134 residues. Highlighted is TSS2, defined by two CAGE peaks that encode truncated ASH2L lacking the first 94 residues corresponding to the IDR region. **(B)** CAGE data across different cell types for human *ASH2L* TSS1 and TSS2. **(C)** RNA-seq data representing the oocyte (Oo), zygote (Z), 2-cell (2C), 4-cell (4C), 8-cell (8C), and morula (M) stages. Shown are fold changes (FC) in normalized read counts relative to the oocyte (Oo) stage for *Macaca mulatta*, *Capra hircus*, and *Sus scrofa*.

**Figure S5. *Ash2l* TSS1 (truncated ASH2L) isoform targets promoters for H3K4me3 important for early development**

**(A)** Western blot of histone H3 lysine 4 methylation (mono, di, and trimethylation) in CRISPRi mESCs or NIH/3T3 fibroblasts targeting TSS1 or TSS2 (left) or cells overexpressing TSS1 or TSS2 ASH2L protein isoforms (right). Blots were also probed for ASH2L, α−Tubulin, and histone H3. A representative blot is shown of n=3 biological repeats. **(B)** Western blot of ASH2L following immunoprecipitation from protein extracts of mESCs: control and CRISPRi targeting TSS1 or TSS2. For the IP, ASH2L antibodies coupled to beads were used. Additionally, extracts of the control were incubated with IgG-coupled beads, which served as a negative control. Highlighted are the full-length and truncated ASH2L protein isoforms. **(C)** Metagene profiles of ASH2L ChIP-seq at protein-coding genes in control mESCs and CRISPRi targeting TSS1 or TSS2. The signals were centred on TSS. ChIP-seq was normalised with chromatin from external *Drosophila melanogaster* (Dm) spike-ins. **(D)** *Ash2l* locus visualized using IGV showing H3K4me3 ChIP-seq signal in mESCs with CRISPRi targeting TSS1 or TSS2. A representative experiment of n=3 biological repeats is shown. **(E)** Metagene profiles and heatmaps of H3K4me3 at protein-coding gene promoters centred on the TSS in CRISPRi mESCs (control, TSS1 or TSS2). **(F)** ChIP-seq of H3K4me3 comparing control to CRISPRi targeting TSS1 or TSS2 in NIH/3T3 cells. Shown is the number of promoters with significant changes (p < 0.01, fold change > 1.3). **(G)** Metagene profiles of H3K4me3 and histone H3 lysine 27 trimethylation (H3K27me3) at gene promoters centred around the TSS in mESCs (Aljazi et al. 2020). Shown are the 575 gene promoters (that showed a significant H3K4me3 difference in the CRISPRi comparing TSS1 and TSS2 in mESCs, see Figure 5F), a random control set of gene promoters, and all protein-coding gene promoters.

**Figure S6. *Ash2* TSS1 (truncated ASH2L) is important for early development.**

**(A)** Expression of pluripotency markers *Oct4*, *Nanog*, and *Klf2*. CRISPRi mESCs were induced to exit pluripotency by culturing in 2i/LIF withdrawal conditions determined by RT-PCR. Data represent the mean ± SD of n = 3 biological replicates. **(B)** Schematic for exit from pluripotency (left). In short, mESCs were gradually withdrawn from 2i/LIF, as described previously(Kalkan et al. 2017). The time points for sample collection are shown. RT-qPCR (right) showing TSS1 and TSS2 *Ash2l* isoform expression levels in CRISPRi mESCs (TSS1 or TSS2) upon 2i withdrawal. **(C)** Gastruloid assays at 96 hours using mESCs with CRISPRi control, or targeting TSS1 or TSS2. Representative images show staining for Brachyury, SOX1, SOX2, and DAPI. **(D)** Quantification of SOX2 and SOX1 distribution in control and CRISPRi mESCs targeting TSS1 or TSS2. Gastruloids were classified based on anterior-posterior (AP) distribution and polarity. Shown are the proportions of gastruloids at 96h, with Brachyury staining polar or away from the pole in combination with SOX2 (control n=74, TSS1 n=34, TSS2 n=27) and SOX1 staining (control n=52, TSS1 n=37, TSS2 n=38). **(E)** Violin plots showing the mean expression levels of E-cadherin, Brachyury, SOX2, and SOX1 along the anterior-posterior axis of elongated gastruloids at 120 hours. Gastruloids were embedded in 5% Matrigel at 96 hours. Also shown is the axial length. A one-way ANOVA, followed by a Dunnett’s multiple comparisons test, was used to compare gastruloids formed from mESCs with CRISPRi (control, TSS1 or TSS2). The samples with statistically significant pair-wise differences are indicated (* for *P*<0.05, ** for *P*<0.01, *** for *P*<0.001, **** for P<0.0001).

**Figure S7. *Ash2l* TSS1 (truncated ASH2L) is important for the differentiation of mESCs into motor neurons.**

**(A)** RT-qPCR quantification of expression of full-length *Ash2l* (TSS2 isoform) and truncated *Ash2l* (TSS1 isoform) during differentiation of mESCs into motor neurons harbouring CRISPRi targeting TSS1 or TSS2. *Gapdh* was used for correcting for relative expression. N = 3 biological replicates; error bars represent SD. **(B)** Western blot of expression of full-length ASH2L (TSS2 isoform) and truncated ASH2L (TSS1 isoform) during differentiation of mESCs into motor neurons harbouring CRISPRi targeting TSS1 or TSS2. α−Tubulin was used as a loading control. A representative blot of n=3 is shown. **(C)** Flow cytometry quantification of SOX2-positive cells (SOX2+), TUBB3/SOX2-negative cells (TUBB3/SOX2-), OLIG2-positive cells (OLIG2+), NKX2.2-positive cells (NKX2.2+), and alive cells in control and CRISPRi targeting TSS1 or TSS2. N = 3 biological replicates; error bars represent SD.

