## Supplementary data for "Lineage-specific retrotransposon co-option reveals a conserved ASH2L isoform as an epigenetic primer of developmental gene promoters"

Figure S1

A

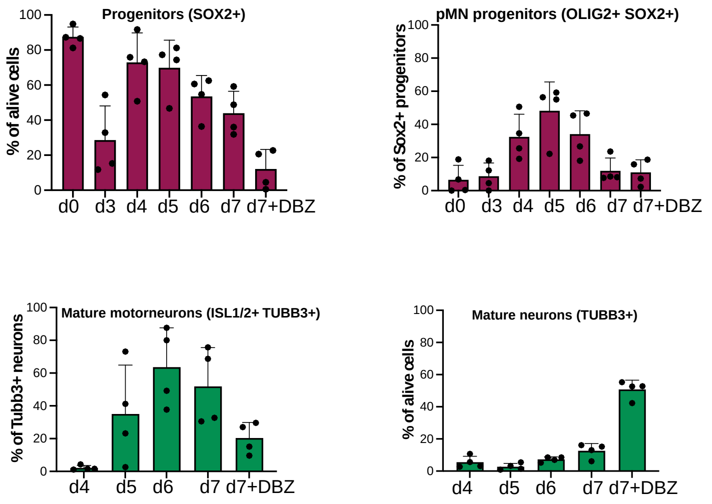

B

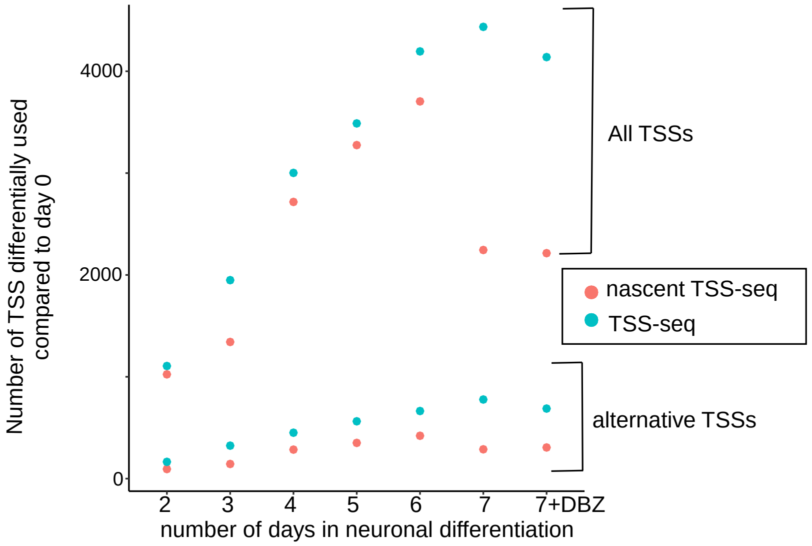

C

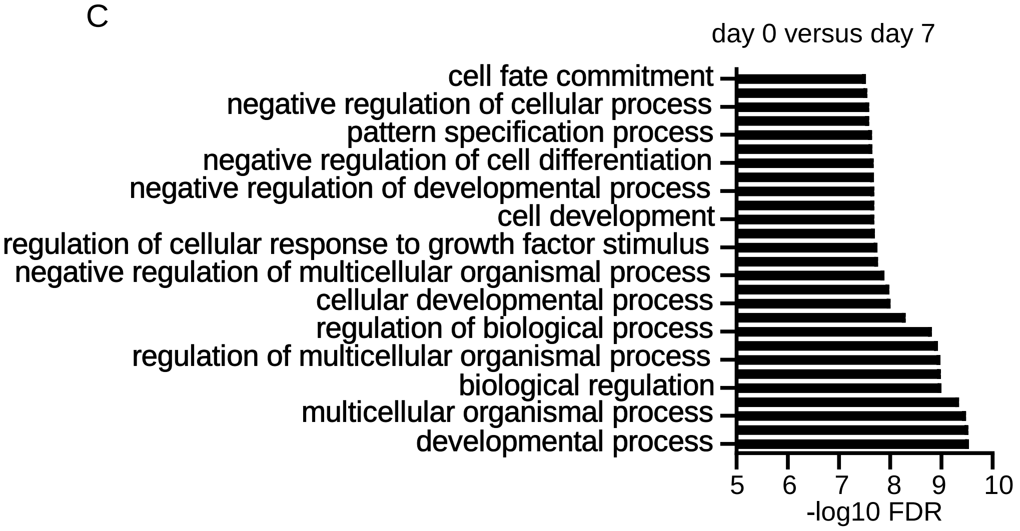

Figure S2

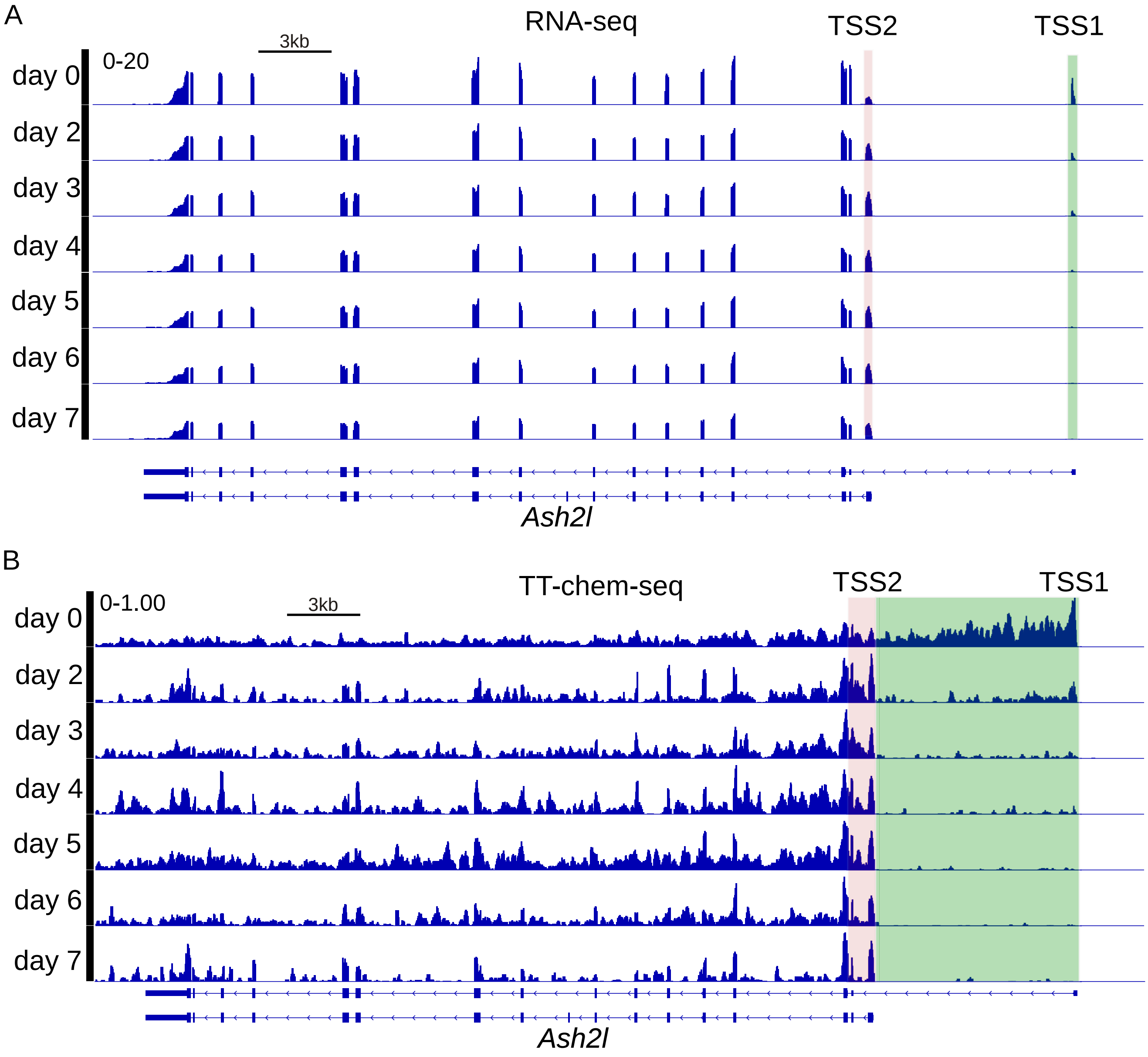

sed

Figure S3

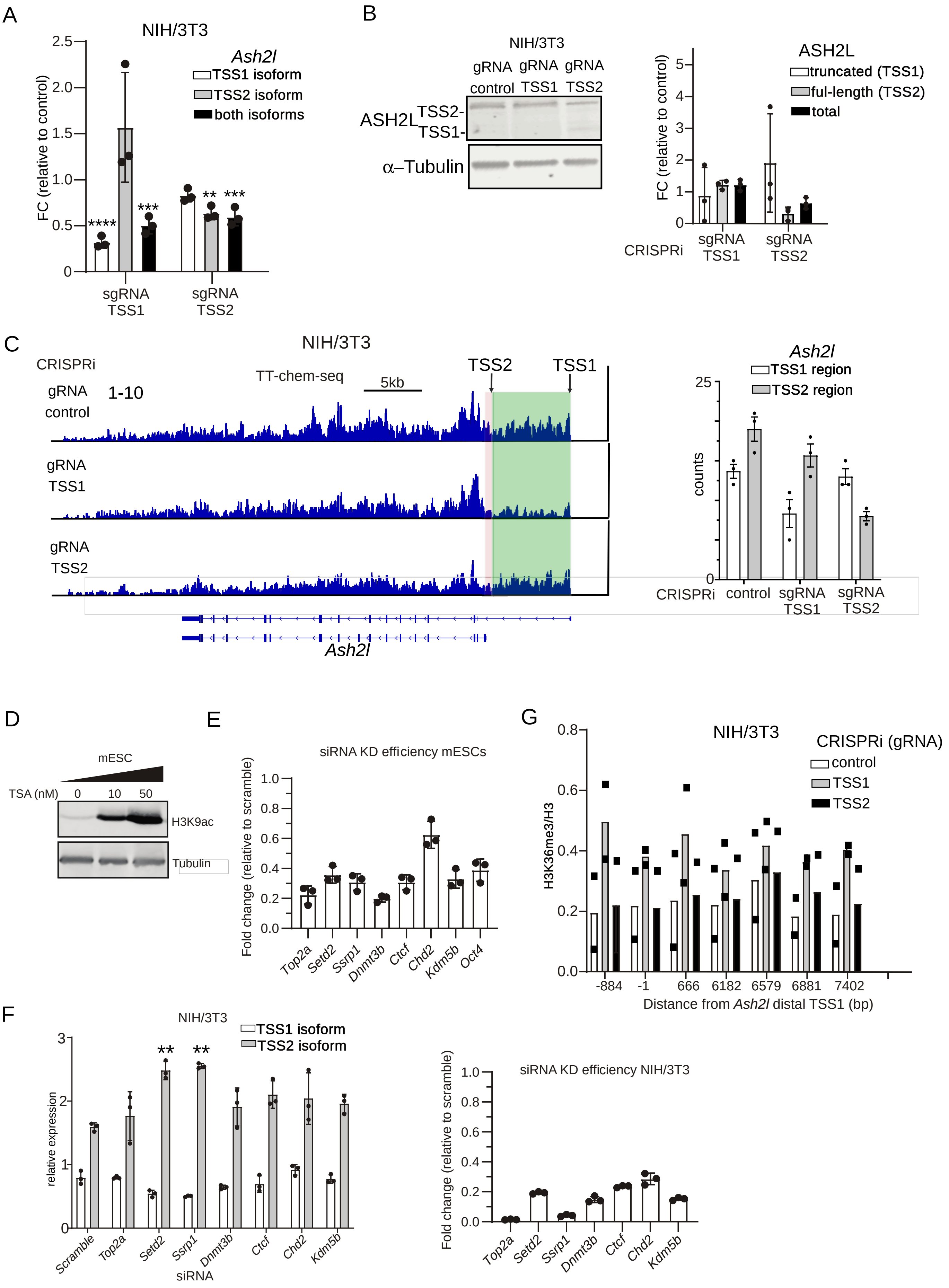

A
NIH/3T3
Ash2l
TSS1 isoform
TSS2 isoform
both isoforms
FC (relative to control)
****
***
**
***
sgRNA TSS1
sgRNA TSS2
B
NIH/3T3
gRNA control
gRNA TSS1
gRNA TSS2
ASH2L
TSS2-
TSS1-
α–Tubulin
ASH2L
truncated (TSS1)
ful-length (TSS2)
total
FC (relative to control)
CRISPRi
sgRNA TSS1
sgRNA TSS2
C
NIH/3T3
CRISPRi
TT-chem-seq
5kb
TSS2
TSS1
gRNA control
1-10
gRNA TSS1
gRNA TSS2
Ash2l
Ash2l
TSS1 region
TSS2 region
counts
CRISPRi
control
sgRNA TSS1
sgRNA TSS2
D
mESC
TSA (nM)
0
10
50
H3K9ac
Tubulin
E
siRNA KD efficiency mESCs
Fold change (relative to scramble)
Top2a
Setd2
Ssrp1
Dnmt3b
Ctcf
Chd2
Kdm5b
Oct4
G
NIH/3T3
CRISPRi (gRNA)
control
TSS1
TSS2
H3K36me3/H3
-884
-1
666
6182
6579
6881
7402
Distance from Ash2l distal TSS1 (bp)
F
NIH/3T3
TSS1 isoform
TSS2 isoform
relative expression
**
**
Scramble
Top2a
Setd2
Ssrp1
Dnmt3b
Ctcf
Chd2
Kdm5b
siRNA
siRNA KD efficiency NIH/3T3
Fold change (relative to scramble)
Top2a
Setd2
Ssrp1
Dnmt3b
Ctcf
Chd2
Kdm5b

Figure S4

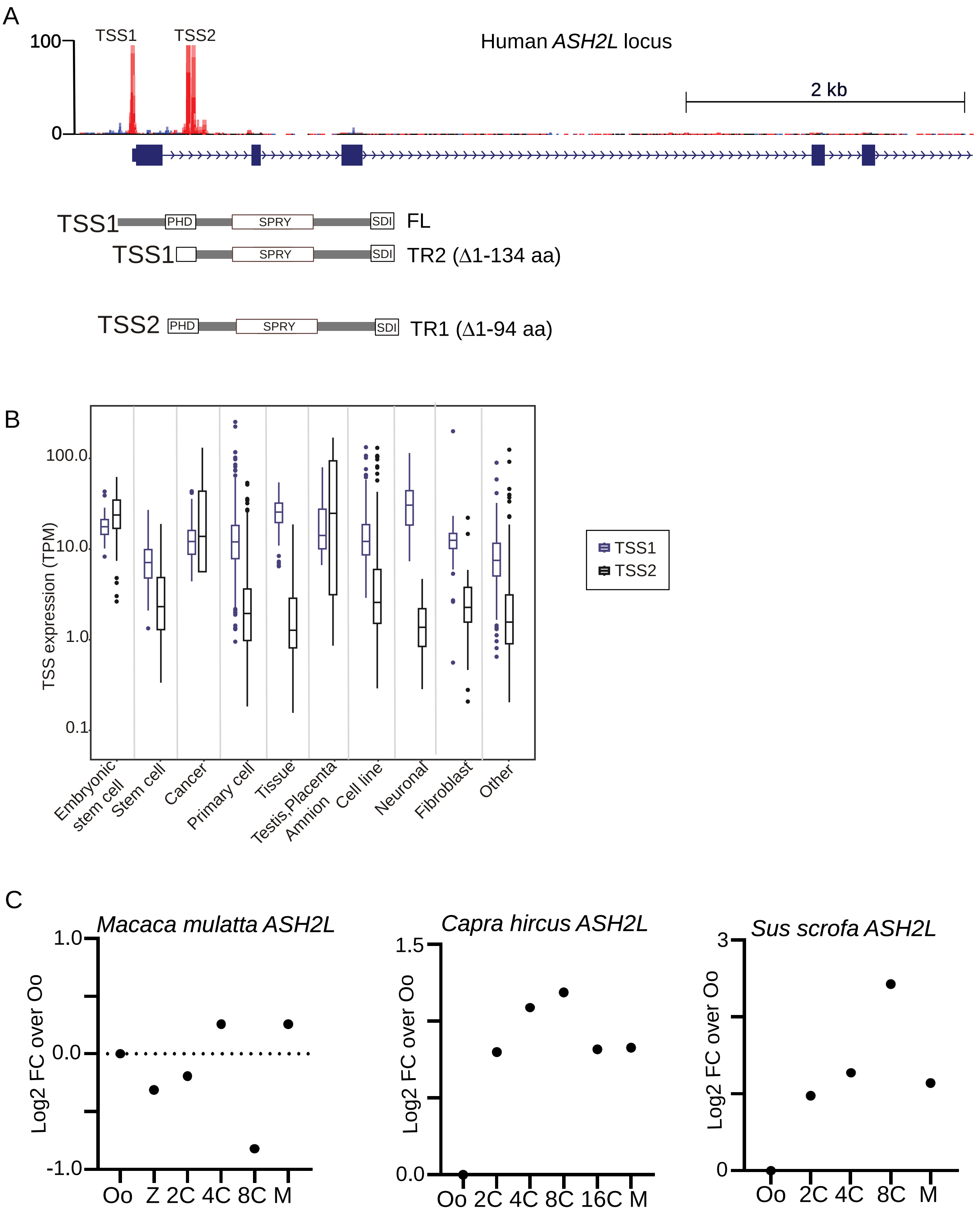

A
Human *ASH2L* locus
TSS1
TSS2
100
0
2 kb
TSS1 PHD SPRY SDI FL
TSS1 SPRY SDI TR2 (Δ1-134 aa)
TSS2 PHD SPRY SDI TR1 (Δ1-94 aa)
B
TSS expression (TPM)
100.0
10.0
1.0
0.1
Embryonic stem cell
Stem cell
Cancer
Primary cell
Tissue
Testis,Placenta Amnion
Cell line
Neuronal
Fibroblast
Other
TSS1
TSS2
C
*Macaca mulatta ASH2L*
Log2 FC over Oo
1.0
0.0
-1.0
Oo Z 2C 4C 8C M
*Capra hircus ASH2L*
Log2 FC over Oo
1.5
0.0
Oo 2C 4C 8C 16C M
*Sus scrofa ASH2L*
Log2 FC over Oo
3
0
Oo 2C 4C 8C M

Figure S5

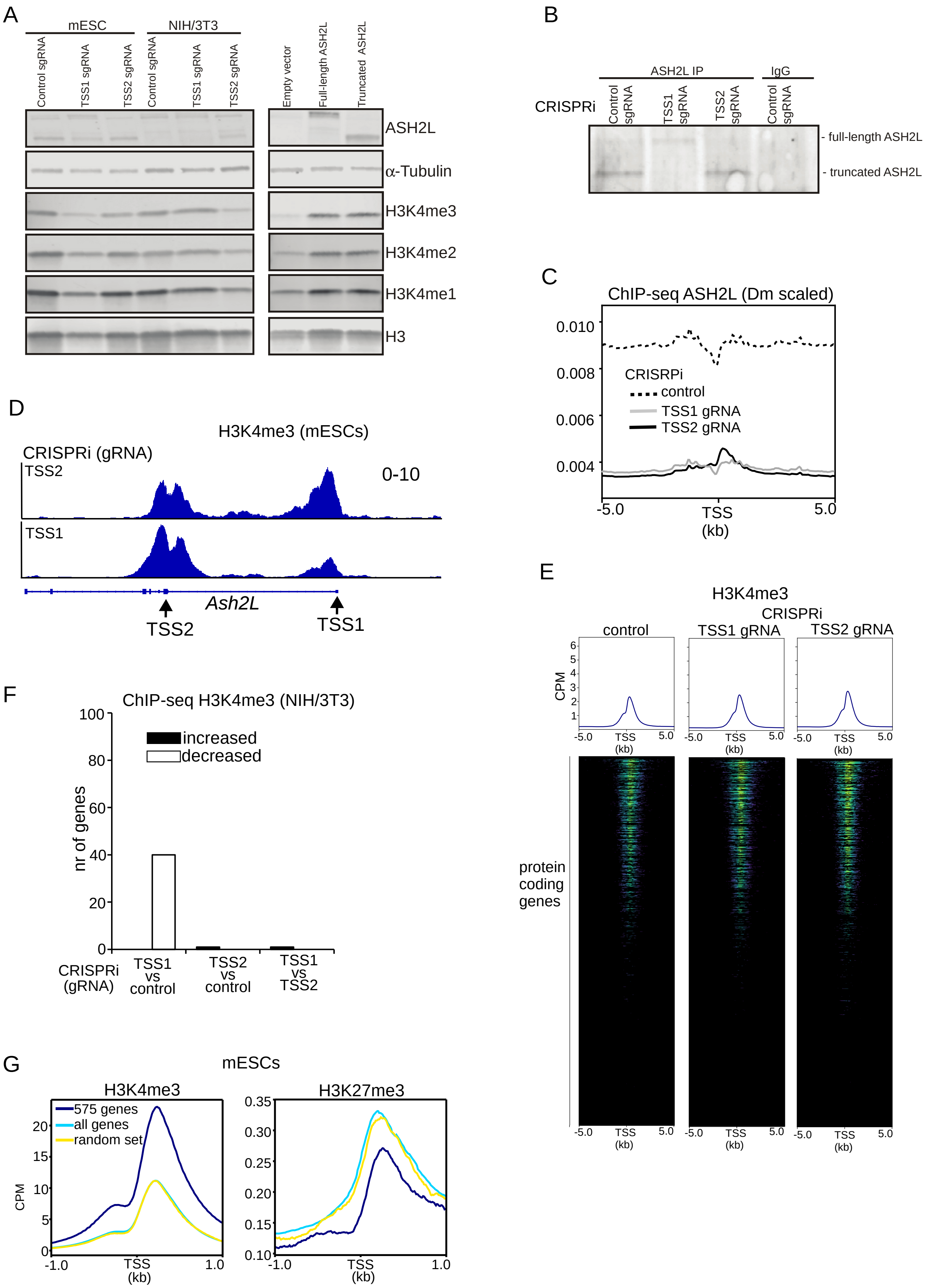

A
mESC
NIH/3T3
Control sgRNA
TSS1 sgRNA
TSS2 sgRNA
Control sgRNA
TSS1 sgRNA
TSS2 sgRNA
Empty vector
Full-length ASH2L
Truncated ASH2L
ASH2L
α-Tubulin
H3K4me3
H3K4me2
H3K4me1
H3
B
ASH2L IP
IgG
CRISPRi
Control sgRNA
TSS1 sgRNA
TSS2 sgRNA
Control sgRNA
- full-length ASH2L
- truncated ASH2L
C
ChIP-seq ASH2L (Dm scaled)
0.010
0.008
0.006
0.004
CRISRPi
control
TSS1 gRNA
TSS2 gRNA
-5.0
TSS
(kb)
5.0
D
H3K4me3 (mESCs)
CRISPRi (gRNA)
TSS2
TSS1
0-10
Ash2L
TSS2
TSS1
E
H3K4me3
CRISPRi
control
TSS1 gRNA
TSS2 gRNA
CPM
6
5
4
3
2
1
-5.0
TSS
(kb)
5.0
protein coding genes
F
ChIP-seq H3K4me3 (NIH/3T3)
100
80
60
40
20
0
nr of genes
increased
decreased
CRISPRi (gRNA)
TSS1 vs control
TSS2 vs control
TSS1 vs TSS2
G
mESCs
H3K4me3
575 genes
all genes
random set
20
15
10
5
0
CPM
-1.0
TSS
(kb)
1.0
H3K27me3
0.35
0.30
0.25
0.20
0.15
0.10
-1.0
TSS
(kb)
1.0

Figure S6

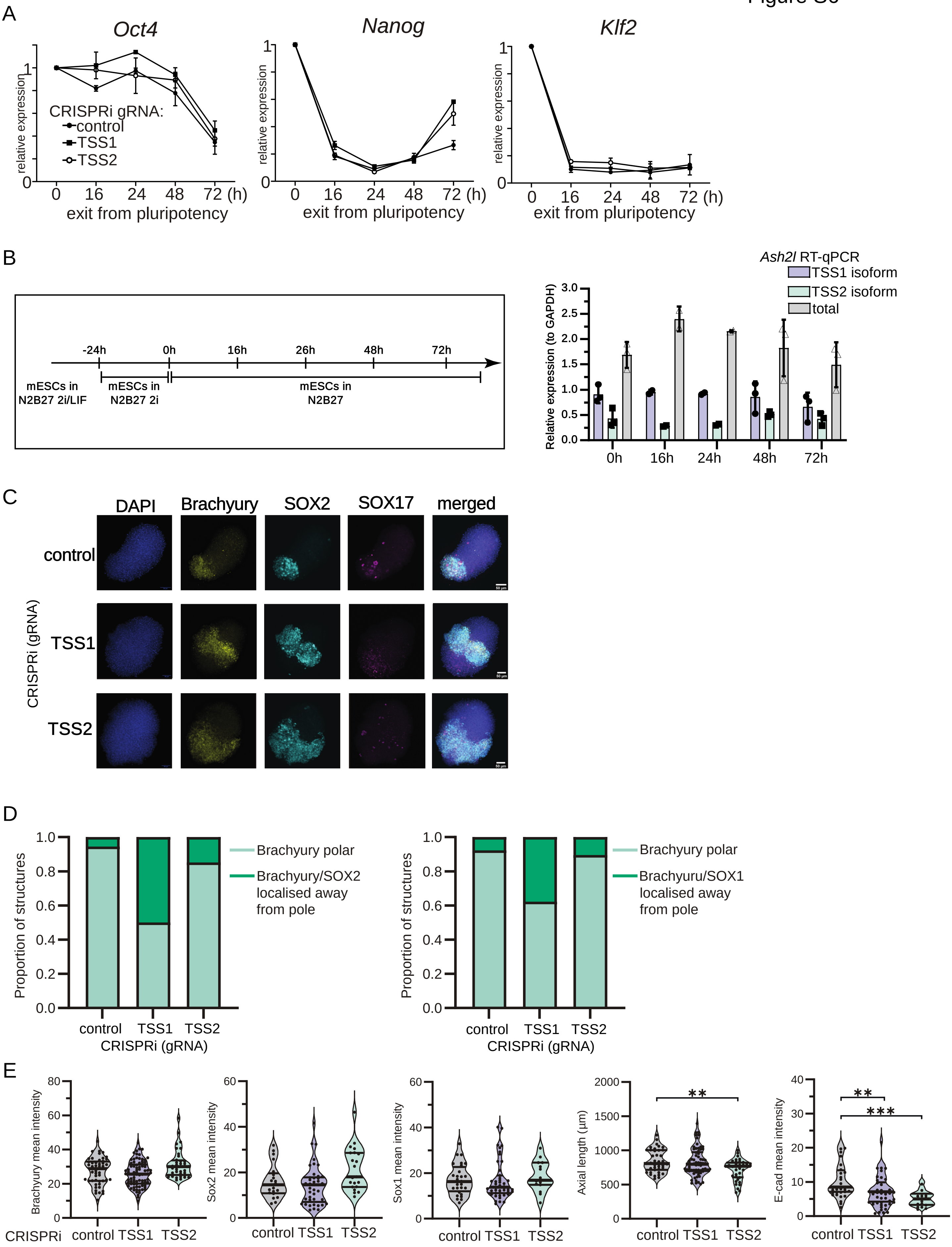
A
Oct4
Nanog
Klf2
relative expression
CRISPRi gRNA:
control
TSS1
TSS2
0 16 24 48 72 (h)
exit from pluripotency
B
-24h 0h 16h 26h 48h 72h
mESCs in N2B27 2i/LIF
mESCs in N2B27 2i
mESCs in N2B27
Ash2l RT-qPCR
TSS1 isoform
TSS2 isoform
total
Relative expression (to GAPDH)
0h 16h 24h 48h 72h
C
DAPI
Brachyury
SOX2
SOX17
merged
control
TSS1
TSS2
CRISPRi (gRNA)
D
Proportion of structures
Brachyury polar
Brachyury/SOX2 localised away from pole
Brachyuru/SOX1 localised away from pole
control TSS1 TSS2
CRISPRi (gRNA)
E
Brachyury mean intensity
Sox2 mean intensity
Sox1 mean intensity
Axial length (µm)
E-cad mean intensity
CRISPRi
control TSS1 TSS2

Figure S7

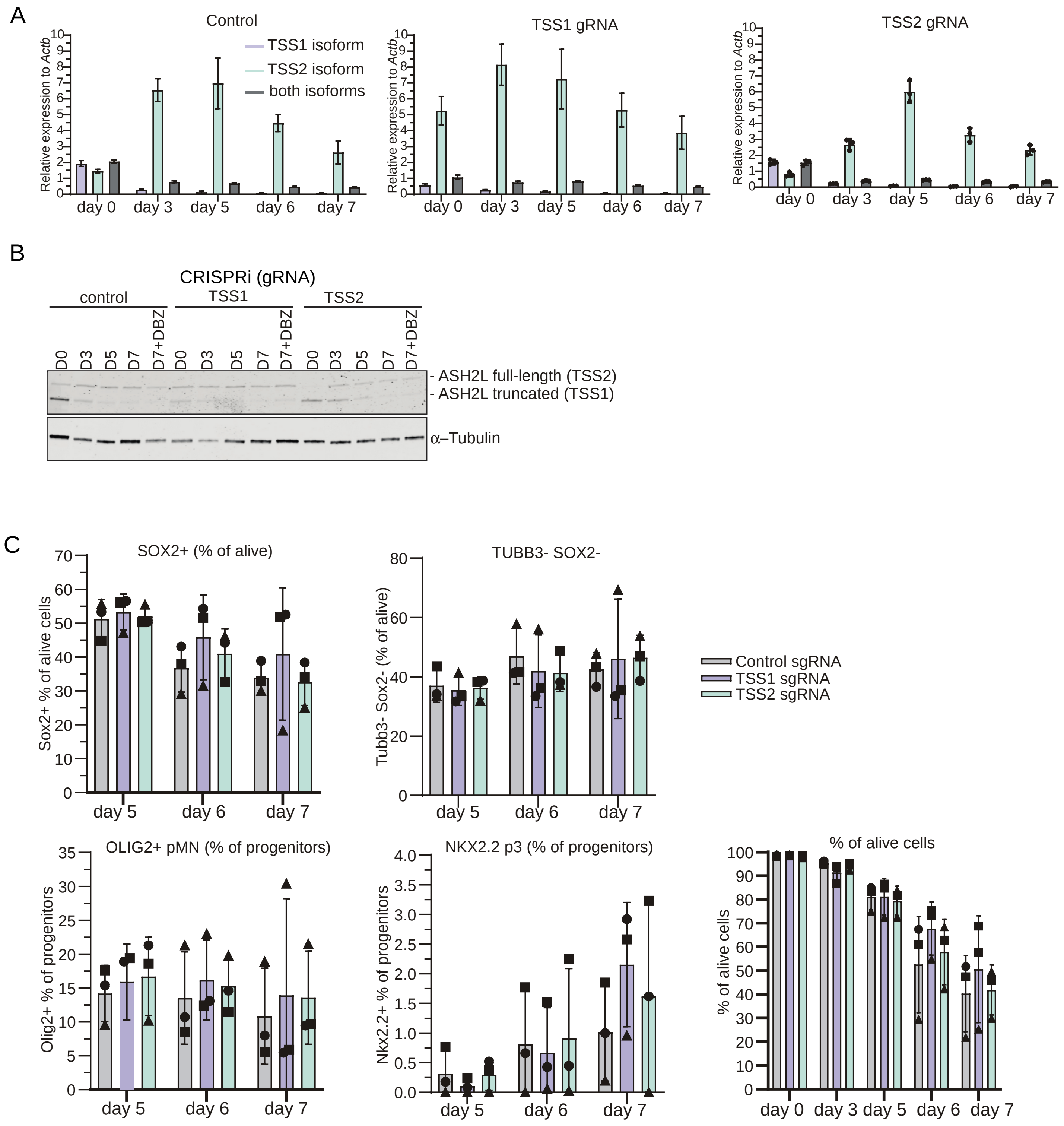
A
Control
TSS1 isoform
TSS2 isoform
both isoforms
Relative expression to Actb
day 0
day 3
day 5
day 6
day 7
TSS1 gRNA
TSS2 gRNA
B
CRISPRi (gRNA)
control
TSS1
TSS2
D0
D3
D5
D7
D7+DBZ
- ASH2L full-length (TSS2)
- ASH2L truncated (TSS1)
α–Tubulin
C
SOX2+ (% of alive)
Sox2+ % of alive cells
TUBB3- SOX2-
Tubb3- Sox2- (% of alive)
Control sgRNA
TSS1 sgRNA
TSS2 sgRNA
OLIG2+ pMN (% of progenitors)
Olig2+ % of progenitors
NKX2.2 p3 (% of progenitors)
Nkx2.2+ % of progenitors
% of alive cells
% of alive cells

**Table S1. Oligo sequences used.**

| Oligo name | Sequence (5'-3') |
|---|---|
| Actin-exon5-Fw | GAAAATCTGGCACCACACCT |
| Actin-exon6-Rv | TAGCACAGCCTGGATAGCAA |
| GAPDH-Fw | CATGACAACTTTGGCATTGTGG |
| GAPDH-Rv | GGATGCAGGGATGATGTTCTG |
| ASH2L-Differentiatedisoform-Fw | CCGAAAGTGGGGATGCAAAC |
| ASH2L-Differentiatedisoform-Rv | AGTGTCTCCGGTACCTTCCA |
| ASH2L-ESCisoform-Fw | GGGAGGTCAACTGGAGTAAGTC |
| ASH2L-ESCisoform-Rv | GCCTGGGTATCCATCACTTCTG |
| ASH2L-universal-Fw | CCCTTAGGTAACCTCCAAGC |
| ASH2L-universal-Fw | GTCCGTAGCCAGACGAATAG |
| Nanog-exon1-Fw | CAGTTTTTCATCCCGAGAAC |
| Nanog-exon2-Rv | CTTTTGTTTGGGACTGGTAG |
| Pou5f1-5UTR-Fw | GTCCCTAGGTGAGCCGTCTTT |
| Pou5f1-exon1-Rv | AGTCTGAAGCCAGGTGTCCAG |
| Sox2-exon1-Fw | GGCAGCTACAGCATGATGCAGGAGC |
| Sox2-exon1-Rv | CTGGTCATGGAGTTGTACTGCAGG |
| Fgf5-Fw | AGAGTGGGCATCGGTTTCCATC |
| Fgf5-Rv | CCTACAATCCCCTGAGACACAG |
| Gata6-Fw | CTAGCGCTGTTTGTTTAGGG |
| Gata6-Rv | TTGATTCCTCGAGCGATGTG |
| Msx1-Fw | GAGGCCAAAAGGACTAGAGG |
| Msx1-Rv | GCGCTTAGAGTTTGACAGTG |
| Ctnnb1-Fw | TGAGACTGCAGATCTTGGAC |
| Ctnnb1-Rv | TCAACTGGATAGTCAGCACC |
| Ccne1-Fw | TCTACTTGGCACAGGACTTC |
| Ccne1-Rv | TAGACATTCAGCCAGGACAC |
| DNMT3B-Fw | GTCAGTACCCCATCAGTTG |
| DNMT3B-RV | CACACGAGGTCACCTATTC |
| SSRP1-Fw | GTACCACAGGCAAGAATGAG |
| SSRP1-Rv | ATCCGGATATCGTATCGACC |
| SETD2-Fw | CTCAAGGTGAAGTAGCATGTGG |
| SETD2-Rv | CTCTCCACAGTATTCCAGGAC |
| KDM5B-Fw | CATCATGATCGAGGACGAG |
| KDM5B-Rv | GTAGGCCAGGTTTACAAGAG |
| TOP2A-Fw | GAGAAGAAAGTGACAGGTGG |
| TOP2A-Rv | CCATGTTATCCATCCACGTC |
| CTCF-Fw | GAGTGGTACCATGAAGATGC |
| CTCF-Rv | GCTGGATGAGAGCATATCG |
| CHD2-Fw | CTGAGTATCTGTGCAAGTGG |
| CHD2-Rv | CCTCCTAAATAAGCTGGCTA |
| EEH_800up_distalTSS_ASH2L_Fw | TCAGACATGAGGAACACACG |
| EEH_distalTSS_A_ASH2L_Fw | GGACTTCCGATTCCAGTGAG |
| EEH_distalTSS_B_ASH2L_Fw | caGTGAGGCAGGTTCGGG |
| EEH_distalTSS_C_ASH2L_Fw | GGAGGTCAACTGGAGTAAGTC |
| EEH_400down_distalTSS_ASH2L_Fw | GGGAAGATATCAAGGGCTGC |

| | |
|---|---|
| EEH_800down_distalTSS_ASH2L_Fw | AACCCGAGGATCTGGGTTTA |
| EEH_700up_proximalTSS_ASH2L_Fw | ACCCATATGCTTGCACTCAG |
| EEH_200up_proximalTSS_ASH2L_Fw | CCAGAGTGCTCCTTTGTTGT |
| EEH_proximalTSS_A_ASH2L_Fw | TTTCCCACTTTAAGAGCGCC |
| EEH_proximalTSS_B_ASH2L_Fw | GCGTCCTTCCCAAGTAAGTG |
| EEH_200down_proximalTSS_ASH2L_Fw | CCGAAAGTGGGTAAGGCAAT |
| EEH_400down_proximalTSS_ASH2L_Fw | CTGATCCCTCTTGCAATGCT |
| EEH_ASH2L_sharedexon3_A_Fw | CTGTGGATGAGGAGAATGGG |
| EEH_ASH2L_sharedexon3_B_Fw | CAGAAGTGATGGATACCCAGG |
| EEH_800downgene_ASH2L_Fw | GGATGCCTTAATGACCTCCG |
| EEH_ActB_TSS_Fw | CCGCTGTGGCGTCCTATAAA |
| EEH_ActB_intron3_Fw | TTCTTGCACTCCTTGCATGT |
| EEH_ActB_highH3K4me3_Fw | CTAGTGTGTCCCCAAGCC |
| EEH_800up_distalTSS_ASH2L_R | GAGCTCCTCCCACTTAGACA |
| EEH_distalTSS_A_ASH2L_R | TTCATCCGTGTCTCAATGGC |
| EEH_distalTSS_B_ASH2L_R | TCCTCCTTCATCCGTGTCTC |
| EEH_distalTSS_C_ASH2L_R | TTGGTCTCGTCCTGGCCT |
| EEH_400down_distalTSS_ASH2L_R | CCTTGACCTGCTGAGTCATT |
| EEH_800down_distalTSS_ASH2L_R | TGTTGGCTGGATCTTGGTTT |
| EEH_700up_proximalTSS_ASH2L_R | GAATTCAGCAAGGTTTGGGC |
| EEH_200up_proximalTSS_ASH2L_R | TTGAAGGGAGCCACTTAGGA |
| EEH_proximalTSS_A_ASH2L_R | ATTGCCTTACCCACTTTCGG |
| EEH_proximalTSS_B_ASH2L_R | CTCTGTCTCTCGGTCCTCC |
| EEH_200down_proximalTSS_ASH2L_R | AGCATTGCAAGAGGGATCAG |
| EEH_400down_proximalTSS_ASH2L_R | ATGCCCACGAGAGAACAATC |
| EEH_ASH2L_sharedexon3_A_R | TATTCCAAAGGTGTCAGCGG |
| EEH_ASH2L_sharedexon3_B_R | CACTGCAACTCCACTTCTCC |
| EEH_800downgene_ASH2L_r | TCTGCATCCTACCTAGCTCC |
| EEH_ActB_TSS_R | AGGAGCTGCAAAGAAGCTGT |
| EEH_ActB_intron3_R | AGCCACAAGAAACACTCAGG |
| EEH_ActB_highH3K4me3_R | TTACGCCTAGCGTGTAGACT |
| EEH_proximalTSS_C_ASH2L_Fw | GCGTCCTTCCCAAGTAAG |
| EEH_proximalTSS_C_ASH2L_R | ATATCTATGCGGCGAGTTG |
| ASH2L_TSS1 gRNA top | ttgGaggagtcgccattgagacagtttaagagc |
| ASH2L_TSS1 gRNA bottom | ttagctcttaaactgtctcaatggcgactcctCcaacaag |
| ASH2L_TSS2 gRNA top | ttgGTGGGAAGGAGCAGCGATGGgtttaagagc |
| ASH2L_TSS2 gRNA bottom | ttagctcttaaacCCATCGCTGCTCCTTCCCACcaacaag |

Table S2. Plasmid used

| Function | Name | Citation |
|---|---|---|
| dCAS9-KRAB | pMH0006 | (Chen et al. 2019) |
| gRNA for EGFP | pCRISPRia-v2-control | (Horlbeck et al. 2016) |
| gRNA TSS1 | pCRISPRia-v2-TSS1 | |
| gRNA TSS2 | pCRISPRia-v2-TSS2 | |
| overexpression ASH2L FL | pCDNA 4/TO-ASH2L-FL | |
| overexpression ASH2L TR | pCDNA 4/TO-ASH2L-TR | |

Chen, John J., Diane L. Nathaniel, Preethi Raghavan, Maxine Nelson, Ruilin Tian, Eric Tse, Jason Y. Hong, et al. 2019. “Compromised Function of the ESCRT Pathway Promotes Endolysosomal Escape of Tau Seeds and Propagation of Tau Aggregation.” *The Journal of Biological Chemistry* 294 (50): 18952–66.

Horlbeck, M. A., L. A. Gilbert, J. E. Villalta, B. Adamson, R. A. Pak, Y. Chen, A. P. Fields, et al. 2016. “Compact and Highly Active Next-Generation Libraries for CRISPR-Mediated Gene Repression and Activation.” *ELife* 5. https://doi.org/10.7554/eLife.19760.

**Table S3. Antibodies used.**

**Western Blot**

| Antigen | Host | Source | Catalog# | Dilution |
|---|---|---|---|---|
| Alpha-Tubulin | Mouse (monoclonal) | Merck | 05-829 | 1:10,000 |
| Histone H3 acetyl K9 | Rabbit (polyclonal) | Abcam | Ab4441 | 1:10,000 |
| H3 | Rabbit (polyclonal) | Abcam | ab1791 | 1:10,000 |
| H3 (2D10) | Mouse (monoclonal) | Abbkine | *ABL1070-50* | 1:10,000 |
| CTCF | Rabbit (polyclonal) | Millipore | 07-729 | 1:1000 |
| ASH2L (D93F6) | Rabbit (monoclonal) | Cell signalling | 5019T | 1:2000 |
| ASH2L | Rabbit (polyclonal) | Active Motif | AB_2615057 | 1:2000 |
| TMEM39A | Rabbit (polyclonal) | Aviva Systems Biology | ABIN2500427 | 1:1000 |
| KDM3A | Rabbit (polyclonal) | Bethyl Laboratories | A301-539A-T | 1:2000 |
| HMGXB4 | Rabbit (polyclonal) | Bethyl Laboratories | A305-022A-T | 1:1000 |
| V5 | Mouse (monoclonal) | Invitrogen | AB_2556564 | 1:2000 |
| Hexokinase | Rabbit (polyclonal) | Stratech | H2035 | 1:8000 |
| Anti-mouse IgG, HRP-linked | Sheep | GE Life Sciences | NA931V5 | 1:10000 |
| Anti-rabbit IgG, HRP-linked | Donkey | GE Life Sciences | NA934V | 1:10000 |
| IRDye 800CW anti-mouse | Goat | LiCOR | 926-32210 | 1:15,000 |
| IRDye 680RD anti-Rabbit | Goat | LiCOR | 926-68071 | 1:15,000 |
| H3K4me3 | Rabbit | Abcam | ab8580 | 1:5000 |
| H3K4me2 | Rabbit | Abcam | ab32356 | 1:5000 |
| H3K4me1 | Rabbit | Abcam | AB8895 | 1:5000 |

**Gastruloid assay**

| Antigen | Dilution | Supplier | Species |
|---|---|---|---|
| Brachyury | 1:150 | Thermo Fisher | Rabbit |
| Sox2 | 1:200 | Thermo Fisher | Rat |
| Sox17 | 1:100 | R&D Systems | Goat |
| Sox1 | 1:100 | R&D Systems | Goat |
| E-Cadherin | 1:200 | Thermo Fisher | Rat |
| Anti-Goat Alexa 488 | 1:200 | Thermo Fisher | Donkey |
| Anti-Rat Alexa 555 | 1:200 | Thermo Fisher | Donkey |

| | | | |
|---|---|---|---|
| Anti-Rabbit Alexa 647 | 1:200 | Thermo Fisher | Donkey |
| Anti-Rat Alexa 488 | 1:200 | Thermo Fisher | Donkey |
| Anti-Goat Alexa 555 | 1:200 | Thermo Fisher | Donkey |

**Motor neuron differentiation**

| Antigen | Dilution | Secondary Antibody | Fluorophore | Supplier | Species |
|---|---|---|---|---|---|
| Goat Olig2 unconjugated | 1:400 | AF488 1:1000 | AF488 | R&D | Goat |
| Nkx2.2-PE (74.5A5) | 1:100 | N/A | PE | BD | Mouse |
| Sox2-PerCPCy5.5 | 1:100 | N/A | PerCP5.5 | BD | Mouse |
| Tubb3-A647 (TUJ1) | 1:100 | N/A | AF647 | BD | Mouse |
| Live/Dead Near IR | 1:1000 | N/A | R-780/60 | Thermo Fisher | N/A |
